# Hydrogen peroxide formation by Nox4 limits malignant transformation

**DOI:** 10.1101/177055

**Authors:** Valeska Helfinger, Florian Freiherr von Gall, Nina Henke, Michael M. Kunze, Tobias Schmid, Juliana Heidler, Ilka Wittig, Heinfried H. Radeke, Viola Marschall, Karen Anderson, Ajay M. Shah, Simone Fulda, Bernhard Brüne, Ralf P. Brandes, Katrin Schröder

**Affiliations:** Institute for Cardiovascular Physiology, Goethe-University, Frankfurt am Main, Germany; Institute for Biochemistry I/Pathobiochemistry, Goethe-University, Frankfurt am Main, Germany; Functional Proteomics, SFB 815 Core Unit, Goethe-University, Frankfurt am Main, Germany; Pharmazentrum Frankfurt, Goethe-University, Frankfurt am Main, Germany; Institute for Experimental Cancer Research in Pediatrics,Goethe-University, Frankfurt, Germany; German Cancer Consortium (DKTK), Heidelberg, Germany, German Cancer Research Center (DKFZ), Heidelberg, Germany; The Babraham Institute, Babraham Research Campus, Cambridge, UK; King’s College London British Heart Foundation Centre, Cardiovascular Division, London, UK; Cluster of Excellence “Macromolecular Complexes”, Goethe University, Frankfurt am Main, Germany

**Keywords:** Nox4, cancer, genomic instability

## Abstract

Reactive oxygen species (ROS) can cause cellular damage and are thought to promote cancer-development. Nevertheless, under physiological conditions, all cells constantly produce ROS, either as chemical by-products or for signaling purpose. During differentiation cells induce the NADPH oxidase Nox4, which constitutively produces low amounts of H_2_O_2_. We infer that this constitutive H_2_O_2_ is unlikely to be carcinogenic and may rather maintain basal activity of cellular surveillance systems.

Utilizing two different murine tumor models we demonstrate that Nox4 prevents malignant transformation and facilitated the recognition of DNA-damage. Upon DNA-damage repair is initiated as consequence of phosphorylation of H2AX (γH2AX). Repair only occurs if nuclear activity of the γH2AX-dephosphorylating phosphatase PP2A is kept sufficiently low, a task fulfilled by Nox4: Nox4 continuously oxidizes AKT, and once oxidized AKT captures PP2A in the cytosol. Absence of Nox4 facilitates nuclear PP2A translocation and dephosphorylation of γH2AX. Simultaneously the proportion of active, phosphorylated AKT is increased. Thus, DNA-damage is not recognized and the increase in AKT activity promotes proliferation. The combination of both events resulted in genomic instability and tumor initiation.

With the identification of the first cancer-protective source of reactive oxygen species, Nox4, the paradigm of reactive-oxygen species-induced initiation of malignancies should be revised.

**Significance:** The stereotype of ROS produced by NADPH oxidases as cause of malignant diseases persists generalized since decades. We demonstrate that the NADPH oxidase Nox4, as constitutive source of ROS, prevents malignant transformation and that its pharmacological inhibition as currently aspired by several companies will potentially increase the risk of malignant cell transformation and eventually tumor formation.

**Precis:** By oxidizing AKT and keeping PP2A in the cytosol, the NADPH oxidase Nox4 allows proper DNA damage repair and averts cancer development.

## Introduction

The notion that reactive oxygen species (ROS) are necessary harmful and are the cause of aging and all main diseases is very prevalent. Nevertheless, the prospective randomized clinical trials on antioxidant supplementation have failed to provide protection and in some studies antioxidants even increased the risk of negative outcome (1, 2). Recently, animal experiments demonstrated that some major antioxidants of clinical use, N-acetylcysteine and vitamin E, accelerate spontaneous lung cancer development in mice carrying mutations in K-Ras and B-Raf (3).

A potential explanation for these findings is that ROS are not just a representation of environmental stress or inflammation but that basically all cells produce ROS for signaling purpose. Such kinds of ROS are generated by the Nox family of NADPH oxidases in a highly controlled fashion (4). Nox-dependent ROS-production in general occurs in response to cellular stimulation through increases in intracellular calcium or activation of Rac and protein kinase C (5). There is however one exception, the NADPH oxidases Nox4. This enzyme constitutively produces low amounts of H_2_O_2_ and thus its output is controlled by the expression level of the enzyme. Hypoxia and TGFβ are potent inducers of Nox4 expression (6, 7). Therefore, increased Nox4 expression in disease conditions can be considered a marker of cellular stress.

Interestingly, some tumor suppressor genes, particularly Nrf2, are induced by ROS and facilitate the antioxidant response and cellular protection (8). In previous work we reported that Nox4 maintains the expression of Nrf2 (9). Moreover, Nox4-/- mice exhibit a hyper-inflammatory response in the vasculature which results in cardiovascular dysfunction and accelerated arteriosclerosis (9–13). We and others also found Nox4 to be induced in the course of differentiation and to be essentially involved in the differentiation process of mesenchymal cells (14–16). As de-differentiation and inflammation are well established pre-requisites for malignant transformation we hypothesized that the knockout of Nox4 increases the risk of malignancy development.

Using two different inflammation driven cancer models as well as cell culture studies, we here demonstrate that endogenous Nox4 maintains genomic stability and attenuates malignant transformation. Nox4 mediates its effect through oxidation of AKT, which subsequently promotes cytosolic PP2A sequestration and PP2A-mediated AKT dephosphorylation. Cytosolic trapping of AKT leads to attenuated nuclear PP2A deposition which facilitates DNA damage detection.

## Materials and Methods

### Materials

The following chemicals were used: 3-methylcholanthrene (MCA), azoxymethane (AOM), NaCl, NH_4_Cl, NaHCO_3_ Hank’s BSS without Ca^2+^ and Mg^2+^ and Trypsin-EDTA solution (T3924) from Sigma-Aldrich (Munich, Germany), Dextran sulphate sodium (DSS) (#16011080; MP Biomedicals, Santa Ana, USA), Dulbecco’s PBS (Gibco lifetechnologies, Carlsbad, CA, USA), Hank’s buffer, SYBR green and Na-EDTA from Applichem (Darmstadt, Germany), TRIS (Carl Roth) and fibronectin (Corning Incorporated, Tewksbury, MA, USA). Collagenase Type 2 was purchased from Worthington (Lakewood, NJ, USA). The PI3-Kinase inhibitor Ly294002 (#BML-ST420) was acquired from Enzo Life Science (Lörrach, Germany). The following antibodies were used: Anti-pH3 (06-570), anti-PP2A and anti-phospho-Histone H2A.X (Ser139) from Millipore (Darmstadt, Germany), anti-AKT, anti-phospho-AKT (Ser473), p-p53 (Ser15), ATM, HA and PTEN, Erk, p-Erk, AKT sepharose bead conjugate and IgG mouse sepharose bead conjugate from CellSignaling (Danvers, MA, USA), anti-p53 (FL-393), anti-p110, NQO1, p-ATM (Ser1981), GFP and anti-Topoisomerase 1 (C-15) from Santa Cruz (Dallas, TX, USA), anti-β-actin (AC-15) from Sigma-Aldrich (Munich, Germany) and anti-p85 from BD transduction laboratories (San Jose, CA, USA). Fluorogenic substrates for proteasome activity were purchased from Boston Biochem (Minneapolis, USA). Human recombinant AKT1 (# ab116412) was purchased from Abcam (Cambridge, United Kingdom).

### Animals and animal procedures

All animal experiments were approved by the local governmental authorities (approval number: F28/27 and F28/46) and were performed in accordance with the animal protection guidelines. C57Bl/6J and Nox2y/-mice were purchased from Charles River (Deisenhofen, Germany). Nox4 -/- mice were generated by targeted deletion of the translation initiation site and of exons 1 and 2 of the Nox4 gene (9), backcrossed into C57Bl/6J for more than 10 generations. Tamoxifen-inducible Nox4-/- mice (Nox4flox/flox-ERT2-Cre+/0 mice) were produced by crossing Nox4flox/flox mice (backcrossed more than 10 generations into C57Bl/6J) with Cre-ERT2+/0 mice (9). Genetic deletion of Nox4 in Nox4flox/flox-ERT2-Cre+/0 mice was achieved by oral tamoxifen administration in the chow (LASCRdiet CreActive TAM400, LASvendi, Soest, Germany) on 10 consecutive days. Both Cre positive as well as the Cre negative littermates received tamoxifen. Nox1y/-mice, kindly provided by Karl-Heinz Krause and previously characterized were used for the same experiments (18). Mice were housed in a specified pathogen-free facility with 12/12 hours day and night cycle and free access to water and chow every time.

To induce fibro sarcomas the chemical carcinogen MCA was injected subcutaneously into the right flank of the mice. In case tumors reached a diameter of 1.5 cm or 150 days after MCA-injection mice were sacrificed and if present the tumor tissue was used for cell isolation, immuno-histological and biochemical analysis.

To analyze the short term effect of MCA-injection mice were sacrificed 20 days after MCA injection and skin of injection site was used for histological and biochemical analysis.

Colon carcinomas were induced as follows: A single dose of 10mg/kg body weight AOM was injected intraperitoneally. After 1 week mice were treated for 1 week with 2% DSS in the drinking water followed by two weeks with usual drinking water. This procedure was repeated 2 additional times. Two weeks after the third DSS cycle mice were sacrificed and the colon was used for further analysis.

### Cell culture

Fibro sarcoma cells were isolated using the tumor dissociation kit for mouse and the gentle MACS Dissociator from Miltenyi Biotec (Bergisch-Gladbach, Germany), following the manufactures instructions. Shortly, tumor tissue was homogenized enzymatically, erythrocytes were lysed and eventually cells were cultured in Dulbecco’s Modified Eagle’s Medium (DMEM) + glutaMAX (Gibco, life technologies; Carlsbad, CA, USA) supplemented with 5% fetal calf serum (FCS) and 1% penicillin (50 U/ml) and streptomycin (50 μg/ml) in a humidified atmosphere of 5% CO_2_ at 37°C. Erythrocyte depletion buffer: 155 mM NH_4_Cl, 10 nM NaHCO_3_ and 100nM EDTA in double distilled water, pH=7.4.

For isolation of dermal skin fibroblasts from mice 1 cm^2^ piece of skin was cut into small pieces, transferred into 2.5 ml Collagenase solution (1000 U/ml in Hanks without Ca^2+^/Mg^2+^) and incubated on shaker for 105 minutes at 37°C. Afterwards the solution was re-suspended and transferred through a 70 μm nylon filter. Cells were centrifuged and placed on a fibronectin-coated dish. The primary fibroblasts are cultured in DMEM/F-12 (Gibco, life technologies) supplemented with 10% FCS and 1% Pen/Strep in a humidified atmosphere of 5% CO_2_ at 37°C. Murine lung endothelial cells were isolated and cultured as described by Schröder et al. (9).

### Site-directed mutagenesis and transfection of plasmids

The QuickChange II XL Site-Directed Mutagenesis Kit (#200521, Agilent) was used to make point mutations of AKT1/2/3. Corresponding primers were designed with QuickChange Primer Design Program. Plasmids for mouse AKT1 (#39531), AKT2 (#64832) and AKT3 (#27293) from Addgene were used for converting the two cysteine residues of the specific AKT into serine to create mutants. The following primers were used for the introduction of single amino acid mutations: AKT1 C296S 5’-ATC CCC TCC TTG CTC AGC CCG AAG TCC-3’ and 5’-GGA CTT CGG GCT GAG CAA GGA GGG GAT-3’; AKT1 C310S 5’-CTC CGG CGT TCC GCT GAA TGT CTT CAT AGT GGC-3’ and 5’-GCC ACT ATG AAG ACA TTC AGC GGA ACG CCG GAG-3’; AKT2 C297S 5’-CTG ATG CCC TCT TTG CTC AAG CCA AAG TCA GTG-3’ and 5’-CAC TGA CTT TGG CTT GAG CAA AGA GGG CAT CAG-3’; AKT2 C311S 5’-TAC TCC GGG GTA CCA CTG AAG GTT TTC ATG GTG-3’ and 5’-CAC CAT GAA AAC CTT CAG TGG TAC CCC GGA GTA-3’; AKT3 C293 S 5’-CTG TGA TCC CTT CTT TGC TAA GCC CAA AAT CCG TAA TTT TT-3’ and 5’-AAA AAT TAC GGA TTT TGG GCT TAG CAA AGA AGG GAT CAC AG-3’; AKT3 C307S 5’-GTA CTC TGG TGT GCC ACT GAA TGT CTT CAT GGT AG-3’ and 5’-TAC CCA TGA AGA CAT TCA GTG GCA CAC CAG AGT AC-3’. The double mutants (C296/310, C297/311, C293/307) were constructed using the single mutants as DNA template and primers for C310S/C311S/C307S. The experiments were performed according to the manufacturers protocol and mutations were verified by sequencing. AKT1 S473A/T308A mutant was a gift from Itamar Goren and AKT K179M from Beate Fisslthaler. Transfection of basal AKT plasmids and mutants was performed with Lipofectamine 3000 (#L3000-015) according to the manufacturer’s instructions.

### Proximity ligation assay (PLA)

Analysis was performed as described in the manufacturer’s protocol (Duolink II Fluorescence, OLink, Upsalla, Sweden). Briefly, isolated fibro sarcoma cells were fixed in phosphate buffered formaldehyde solution (4%), permeabilized with Triton X-100 (0.2%), blocked with serum albumin solution (3%) in phosphate buffered saline and incubated overnight with appropriate antibodies. After washing, samples were incubated with respective PLA-probes for 1 hour, washed and ligated for 30 min, both at 37°C. An additional washing step followed and amplification with polymerase was performed for 100 min. Images were obtained by confocal microscopy with LSM 510 (Zeiss, Jena, Germany). Fiji software was used for quantification of single dots per cell.

### Protein and Western Blot analysis

For whole cell protein isolation, cells were lysed in a buffer containing 20mM TRIS/cl pH 7.5, 150 nM NaCl, 10mM NaPP_i_, 20 nM NaF, 1% Triton, 10nM Okadaic acid (OA), 2mM Orthovanadat (OV), protein-inhibitor mix (PIM) and 40 μg/ml phenylmethylsulfonylfluorid (PMSF).

For separation of nucleus and cytosol, the cells were lysed in buffer A (10 nM HEPES pH 7.9, 10 nM KCL, 0.1 mM EDTA, 0.1 mM EGTA, 1% Nonidet, 10 mM DTT, protein-inhibitor mix (PIM), 40 μg/ml phenylmethylsulfonylfluorid (PMSF). Cells were centrifuged to gain the cytosol containing supernatant. The pellet was further lysed with buffer B (20 mM HEPES pH 7.9, 0.4 M NaCl, 1 mM EDTA, 1 mM EGTA, 10 mM DTT, protein-inhibitor mix (PIM), 40 μg/ml phenylmethylsulfonylfluorid (PMSF)) to obtain the nuclear extract. Bradford assay was used to determine the protein amount (19). Samples were cooked in sample buffer and were transferred on SDS-PAGE followed by Western Blotting. Identical amounts of protein from nuclear and cytosolic fractions were loaded. Analysis was performed with an infrared-based detection system using fluorescent-dye-conjugated secondary antibodies from LI-COR biosciences.

Phosphatase activity was analyzed according to the manufacturer’s suggestion. Briefly, 50 mM pNPP were incubated with lysates of cytosolic or nuclear fractions prepared as described above. After 5-10 minutes the reaction was stopped by addition of NaOH and the amount of the product of the phosphatase reaction, p-nitrophenol, was determined by reading the absorbance at 405 nm in a microplate reader (Tecan Infinite 200 Pro).

BIAM Switch Assay was performed to determine the oxidation state of AKT. Briefly, cells were blocked with n-ethylmaleimide (NEM) and scratched in TCA. To block the free thiols cell pellets were resuspended in NEM-denaturing buffer (containing Tris-HCL pH 8.5, Urea, EDTA and SDS) subsequently followed by acetone precipitation. Reduction of oxidized thiols was performed with DTT-denaturing buffer and labeling of those with biotin-polyethyleneoxide-iodoacetamide-denaturing buffer followed by acetone precipitation and triton lysis. Lysates were used for IP with streptavidin-agarose beads and blotted for AKT.

### mRNA isolation and RT-qPCR

Total mRNA from cells and frozen homogenized tissue was isolated with a RNA-Mini-kit (Bio&Sell, Feucht, Germany) according to the manufacturers protocol. Random hexamer primers (Promega, Madison,WI, USA) and Superscript III Reverse Transcriptase (Invitrogen, Darmstadt, Germany) were used for cDNA synthesis. Semi-quantitative real-time PCR was performed with Mx3000P qPCR cycler (Agilent Technologie, Santa Clara, CA, USA) using PCR Eva Green qPCR Mix with ROX (Bio&Sell, Feucht, Germany) with appropriate primers. Relative expression of target genes were normalized to eukaryotic translation elongation factor 2 (EF2), analyzed by delta-delta-Ct method and given as percentage compared to control experiments. Primer sequences for murine Nox4 were forward 5’-TGTTGGGCCTAGGATTGTGTT-3’ and reverse 5’-AGGGACCTTCTGTGATCCTCG-3’. For p53 forward 5’-AGACCGCCGTACAGAAGAAG3’ and reverse 5’-TTCAGCTCCCGGAACATCTC-3’, Cyp1A1 forward 5‘-GGCCACTTTGACCCTTACAA-3’ and reverse 5‘-CAGGTAACGGAGGACAGGAA-3’-3’, for Cyp1B1 forward 5‘-TTCTCCAGCTTTTTGCCTGT-3’ and reverse 5‘-TAATGAAGCCGTCCTTGTCC-3’ and ATM forward 5’-ATGCCAGTCTTTTCAGGGTG-3‘ and reverse 5‘-TCAGAAGCTGGGAGTGCTTC-3’.

### DNA-damage detection (Comet- and Nicoletti-Assay)

A suspension of 1*10^6^ cells/ml was mixes 1:10 with 5% low melting agarose and subjected onto slides coated with 1.5% normal melting agarose. Lysis of the cells was performed for 2 hours at 4°C in lysis buffer (2.5 M NaCl, 10 mM TRIS, 100 mM EDTA, pH=10., 1% Triton X-100 and 10% SDS in double distilled water). Lysis was followed by a 20 minutes incubation of the slides on ice with the alkaline electrophoresis buffer (300 mM NaOH and 0.5 M EDTA) with subsequent electrophoresis at 25 V for 20 minutes. Slides were washed three times with PBS and were stained with SYBR green. Pictures were taken with a confocal microscope LSM 510 Meta and quantification was done manually by three independent investigators determining the ratio of cell number/cells with comets.

Additionally DNA fragmentation was determined by analysis of propidium iodide (PI)-stained nuclei using flow cytometry as described previously (20). Cells were lysed and stained for 2 h in a solution of 0.1% tri-sodium citrate dehydrate and 0.1% Triton X-100 containing 50 μg/ml propidium iodide (Sigma, Deisenhofen, Germany) and analyzed by flow cytometry (FACSCanto II, BD Biosciences, Heidelberg, Germany).

### Immunohistochemistry

Tissue was fixed in 4% paraformaldehyde overnight, dehydrated in ascending ethanol-series and then embedded in paraffin. Paraffin blocks were sliced and slides were dewaxed for further staining in descending ethanol-series from 100% to 70%. For antigen retrieval slides were cooked 10 min in citrate 1 x TRS buffer (Dako). After cooling down staining procedure started using the Dako CSA system in combination with Biotin blocking System (Dako) for 3,3’ diaminobenzidine staining. First, slides were block with Avidin and Biotin for 20 minutes with the biotin blocking system from Dako, followed by peroxidase and protein blocking from the Dako staining kit. In between washing steps were performed with TBST. After the protein block first antibody was applied on the slides overnight at 4°C. The next day slides were incubated with biotinylated secondary antibody, amplification steps were performed as described in the manual and diaminobenzidine was used to generate the signal. Counterstaining was performed with hematoxylin.

### Polysomal Fractionation

Isolated fibro sarcoma cells were seeded in a 15 cm dish one day prior to harvesting. Polysome analysis was performed as described previously (21). Briefly, after incubation with 100 μg/ml cycloheximide for 10 min at 37°C cells were scraped and centrifuged. The supernatant was discarded, pellet washed, centrifuged again and lysed in 750 μl polysome buffer (140 mM KCl, 20 mM Tris-HCl pH 8.0, 5 mM MgCl2, 0.5% NP40, 0.5 mg/mL heparin, 1 mM DTT, 100 U/mL RNasin (Promega), 100 μg/mL CHX). After pelleting for 5 min, 16000 rpm at 4°C and transferring the supernatant into a fresh tube, 600 μl of cytoplasmic lysate was layered onto a 10-50% continuous sucrose gradient. The gradient was centrifuged at 35000 rpm for 2 h without brake, and the gradients were collected in 1-mLfractions using a Gradient Station (Biocomp). Absorbance was measured at 254 nm. RNA was precipitated by addition of sodium acetate (3 M) and isopropanol. RNA was further purified using the Nucleospin RNA Kit (Macherey-Nagel) according to the manufacturer’s manual and analyzed by RT-qPCR.

### Mass spectrometry measurements of inositol lipids

Mass spectrometry was used to measure inositol lipid levels essentially as described previously (22) using a QTRAP 4000 (AB Sciex) mass spectrometer and employing the lipid extraction and derivatization method described for cultured cells (cells isolated from tumors) and whole tissue (tumor tissue), with the modification that initial samples were probe sonicated for 5” (using a microtip) prior to extraction and that final samples were dried in a speedvac concentrator rather than under N2. Measurements were conducted on 1 x10^6^ isolated fibro sarcoma cells or 1mg wet weight tumor tissue per sample. C16:0/C17:0 PI (100ng) and PIP3 (10ng) internal standards (ISDs) were added to each sample prior to extraction. Integrated area of lipid species peaks were corrected for recovery against ISD area (giving a response ratio for each lipid) and data expressed as C18:0 C20:4 PIP3 response ratio normalized to C18:0 C20:4 PI response ratio to account for cell input variation. Both endogenous PIP3 and PI were corrected to their own internal standard, and then the one measurement is divided by the other, to get the best estimate of true PIP3/PI.

### Sample preparation and mass spectrometry

2.5 μM recombinant AKT1 (Abcam) were reduced in 100 μM DTT and treated with 300μM H_2_O_2_ for 30 min. Proteins were directly digested overnight with trypsin (sequencing grade, Promega) and analyzed by liquid chromatography / mass spectrometry (LC/MS).

Fibro sarcoma cells of wildtype and Nox4-deficient mice were lysed in buffer A. Lysates were used to trap AKT1-3 including interacting proteins using a sepharose beads immobilized AKT antibody and IgG (CellSignaling) as negative control. Beads were washed in PBS, resuspended in 50 μl 8 M Urea, 50 mM Tris/HCl, pH 8.5, and incubated at 22°C for 30 min. Thiols were alkylated with 40 mM chloroacetamid and samples were diluted with 25 mM Tris/HCl, pH 8.5, 10% acetonitrile to obtain a final urea concentration of 2 M. Proteins were digested with 1 μg Trypsin/LysC (sequencing grade, Promega) overnight at 22°C under gentle agitation. Digestion was stopped by adding trifluoroacetic acid to a final concentration of 0.1 %. Peptides were loaded and purified on multi-stop-and-go tip (StageTip) containing six C18-disks (23). Peptides were dried and resolved in 1% acetonitrile, 0.1 % formic acid and analyzed by LC/MS.

LC/MS was performed on Thermo Scientific™ Q Exactive Plus equipped with an ultra-high performance liquid chromatography unit (Thermo Scientific Dionex Ultimate 3000) and a Nanospray Flex Ion-Source (Thermo Scientific). Peptides were loaded on a C18 reversed-phase precolumn (Thermo Scientific) followed by separation on a with 2.4 μm Reprosil C18 resin (Dr. Maisch GmbH) in-house packed picotip emitter tip (diameter 100 μm, 15 cm long, New Objectives) using an gradient from mobile phase A (4% acetonitrile, 0.1% formic acid) to 30 % mobile phase B (80% acetonitrile, 0.1% formic acid) for 20 min (purified human AKT1) or 90 min (AKT1 enriched from lysate) followed by a second gradient to 60% B for 10 min or 15 min, respectively. MS data were recorded by data dependent acquisition Top10 method selecting the most abundant precursor ions in positive mode for HCD fragmentation. The full MS scan range was 300 to 2000 m/z with resolution of 70000, and an automatic gain control (AGC) value of 3*10^6^ total ion counts with a maximal ion injection time of 160 ms. MS/MS scans were recorded with a resolution of 17500, an isolation window of 2 m/z and an AGC value set to 10^5^ ions with a maximal ion injection time of 150 ms. Selected ions were excluded in a time frame of 30s following fragmentation event.

Data analysis of purified human AKT1: Xcalibur raw files were analyzed by Peaks7 Studio software for proteomics (www.bioinfor.com; Bioinformatics Solutions, Waterloo, ON, Canada). The enzyme specificity was set to trypsin. Missed cleavages were limited to 3. Monoisotopic precursor mass error tolerance was 5 ppm, and fragment ion tolerance was 0.05 Da. Following variable modifications were selected: of methionine (+15.99), disulfide bridge (-2.02), half of disulfide bridge (-1.01), dioxidation on cysteines (+31.99) and trioxidation on cysteines (+47.98). After de novo sequencing of spectra, the human reference proteome set (download from Uniprot, April 4th, 2015; 68511 entries; www.uniprot.org) was used to identify peptide-spectrum matches with a false discovery rate (FDR) of 1%. For identification of crosslinked peptides by disulfide bridges, a special software StavroX (v3.4.12) (24) was used. Disulfide bridge (-2.01565) was included and the search for dipeptides was done using precise scoring mode. Only di-peptides with highest scores (> 200) were inspected and shown in supplementary table IW2. Disulfide bridges within the structure of AKT1 (4EJN, (25)) were illustrated using Pymol (0.99rev9)).

Data analysis of immune-trapped AKT1-3: Xcalibur raw files were analyzed by MaxQuant (1.5.2.8 (26)) with specificity to trypsin and tolerated missed cleavages of 2. Following variable modifications were selected: of methionine (+15.99), carbamidomethylation on cysteines (+57.02) and acetylation on N-terminus (+42.01). Proteins were identified using proteome set of mouse (Uniprot, 26th June, 2015, 76086 entries) with a false discovery rate (FDR) of 1%. Proteins were quantified using lable free quantification (LFQ) with at least one peptide and further analyzed by Perseus 1.5.4.1. (27). Reverse hits, known contaminants and identified by side proteins were removed from the list. Missing LFQ values were replaced by a background value generated from the normal distribution to enable calculation of ratios to control. Permutation test was performed to identify interacting proteins on AKT1-3.

### Statistics

All values are mean±SEM. Statistical analysis was performed by ANOVA followed by LSD post hoc testing or by t-test, if appropriate. Tumor free survival curves were compared by ANOVA for repeated measurements. Densitometry was performed with the odyssey-software. A p-value of less than 0.05 was considered statistically significant.

## Results

### Loss of Nox4 promotes tumor development

In order to determine the impact of Nox4 on tumor development, knockout mice were used. As compared to WT mice, Nox4 knockout mice develop twice as many tumors in the AOM/DSS-colon carcinoma model (Fig. 1A-C). Similar results were obtained in a second model, fibrosarcoma-development in response to MCA. In Nox4-/- mice, as compared to WT animals, the onset of tumor development was earlier and the developing tumors grew faster (Fig. 1D-F). Proliferation within the both tumor models was more than twice as high in Nox4-/- mice as compared to WT mice (Fig. 1G&H). Importantly, these effects were specific for Nox4: Deletion of Nox1 rather had an inhibitory effect on tumor development, and deletion of Nox2 was without effect at all (Supplemental figure 1). To substantiate the importance of Nox4 as inhibitory enzyme for tumor development, conditional Nox4 knockout mice were studied. Tamoxifen-induced systemic deletion of Nox4 resulted in a similar reduction of tumor development as observed in the global knockout mice. Conditional deletion of Nox4 in macrophages or endothelial cells, in contrast, was without effect on tumor development, suggesting that most probably Nox4 in the transforming cells but not in the tumor stroma mediates the protective effect (Supplemental figure 2).

**Figure 1.**
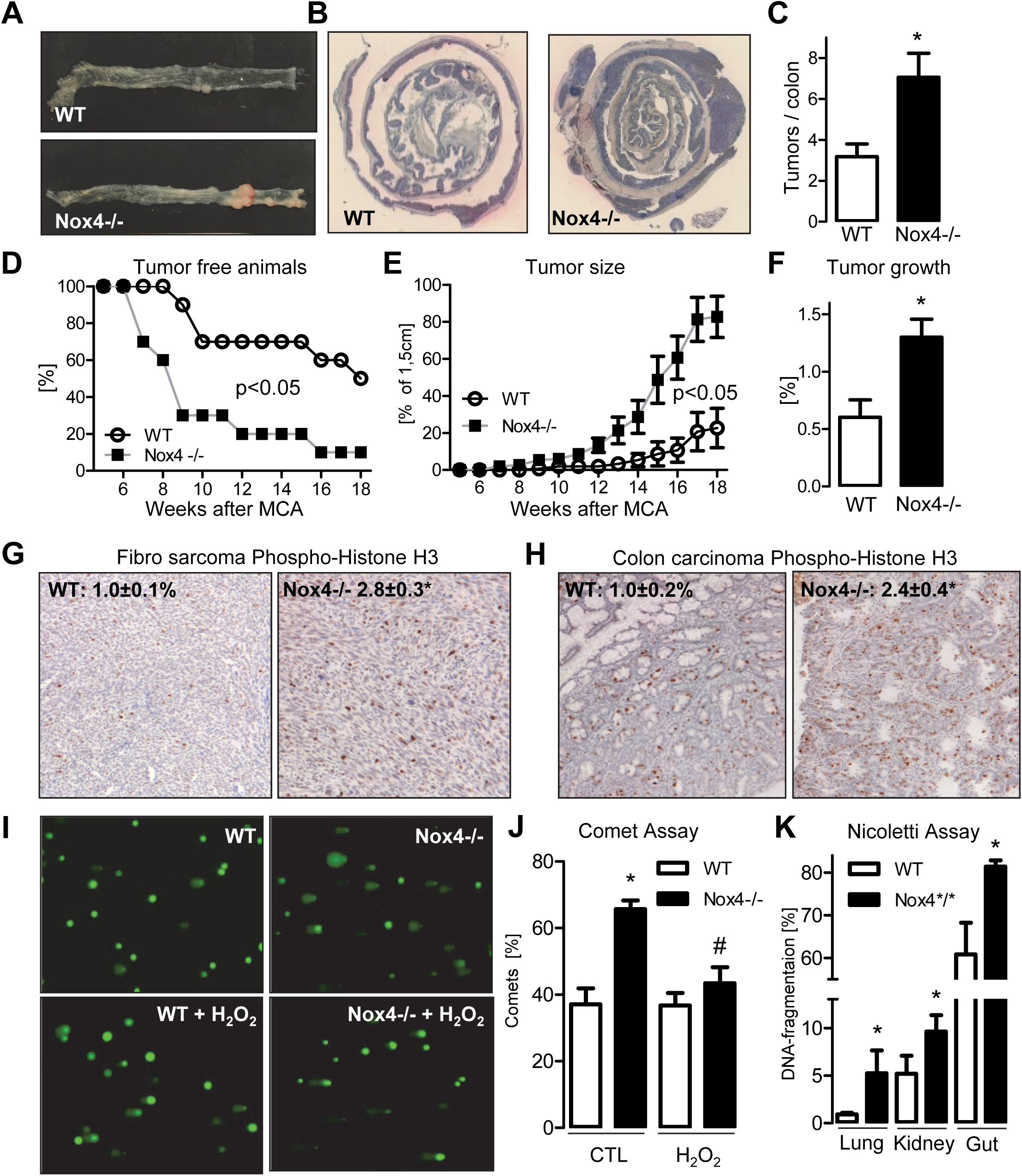
Nox4 knockout promotes tumor development and induces genomic instability. (**A-C**) AOM-DSS colon carcinogenesis model: Representative macro photo (**A**), H&E overview (**B**) and average number of solid carcinoma nodules (**C**) in the colon induced by AOM/DSS. (n=11), *p<0.05 (**D-F**): MCA fibro sarcoma model. Fibro sarcoma development (**D**) and tumor size (**E**) (100% equals the maximal tumor size of 1.5 cm, n=10). (**F**) Relative tumor growth. (**G&H**). Sarcomas and colon carcinomas stained for phosphorylated histone H3 (pH3) as proliferation marker. The statistics indicate the relative staining intensity (n=5-9). (**I&J**) Measurement of DNA double-strand breaks in fibro sarcoma cells was performed using Comet Assay (n=5). Comet Assay of cells treated with or without H_2_O_2_ (24h, 5 μM). Original pictures (**I**) and statistics (**J**) *p<0.05; WT vs. Nox4-/-, #p<0.05 CTL vs. H_2_O_2_ treated. (**K**) Conditional tamoxifen-inducible knockout mice were treated with tamoxifen for 10 day and one week later cells were isolated from multiple organs and analyzed for DNA-fragmentation by Nicoletti Assay (n=3).

### Knockdown of Nox4 results in genomic instability

As genomic instability is an important factor promoting tumor development, cell lines from WT and Nox4 tumors were established and comet assays as a marker for DNA fragmentation were performed. Nox4 knockout tumor cells exhibited a doubling in the number of comets. Importantly, this effect was a consequence of a shortage of H_2_O_2_, the product of Nox4. When the tumor cells were treated with H_2_O_2_ (5μmol/L, 24h) the number of comets in Nox4-/- tumor cells was reduced to that in the WT tumor cells, whereas WT tumor cells did not respond to this low concentration of H_2_O_2_ (Fig. 1I&J). These observations suggest that lack of Nox4-derived H_2_O_2_ may promote genomic instability. Indeed, Nicoletti staining of lung, liver and intestine of healthy Nox4-/- mice demonstrated that DNA strand breaks are more frequent after deletion of Nox4 even in normal tissue (Fig. 1K).

### Loss of Nox4 results in attenuated recognition of DNA damage

The increased DNA fragmentation in healthy Nox4-/- mice points towards an attenuated recognition or DNA damage response after deletion of the ROS generator. Indeed, markers of DNA damage like phosphorylation of histone 2AX (here termed γH2AX) were decreased in tumors and tumor cells of Nox4-/- mice as compared to WT mice. Importantly, already in the healthy colon of Nox4-/- mice or in fibroblasts cultured from those mice, the levels of γH2AX were attenuated, which speaks for a general attenuation of the DNA damage response (Fig. 2A-D). Accordingly, acute exposure of fibroblasts to the carcinogen MCA increased γH2AX less effectively in Nox4-/- cells as compared to WT cells (Fig. 2D). P53 is an important transcription factor responding to DNA damage. In line with the γH2AX data, p53 protein and mRNA expression was reduced in Nox4 knockout tumors and cells isolated from the tumor as compared to WT controls (Fig. 2E-H). Importantly, treatment with H_2_O_2_, the product of Nox4 normalized p53 mRNA expression in Nox4-deficient cells but had no effect in WT cells (Fig. 2F). The decreased p53 expression was not a consequence of altered phosphorylation or translation of p53 (Supplemental figure 3A-D) nor was the expression of enzymes involved in p53 degradation, like NQO1 and MDM2 increased in Nox4-/- cells (Supplemental figure 3E&F). In order to study whether the reduced p53 expression is a consequence of an inappropriate response to damage, we stimulated healthy endothelial cells cultured from the lung of WT and Nox4-/- mice with the carcinogen MCA. Whereas p53 expression was strongly induced in cells of WT mice, the knockout mice exhibited a highly attenuated response (Fig. 2I). This indicates that DNA damage response is attenuated after deletion of Nox4. Importantly, expression and activity of ATM, the kinase which phosphorylates H2AX in response to damage and initiates the DNA damage response was not reduced in Nox4-deficient cells (Supplemental figure 4).

**Figure 2.**
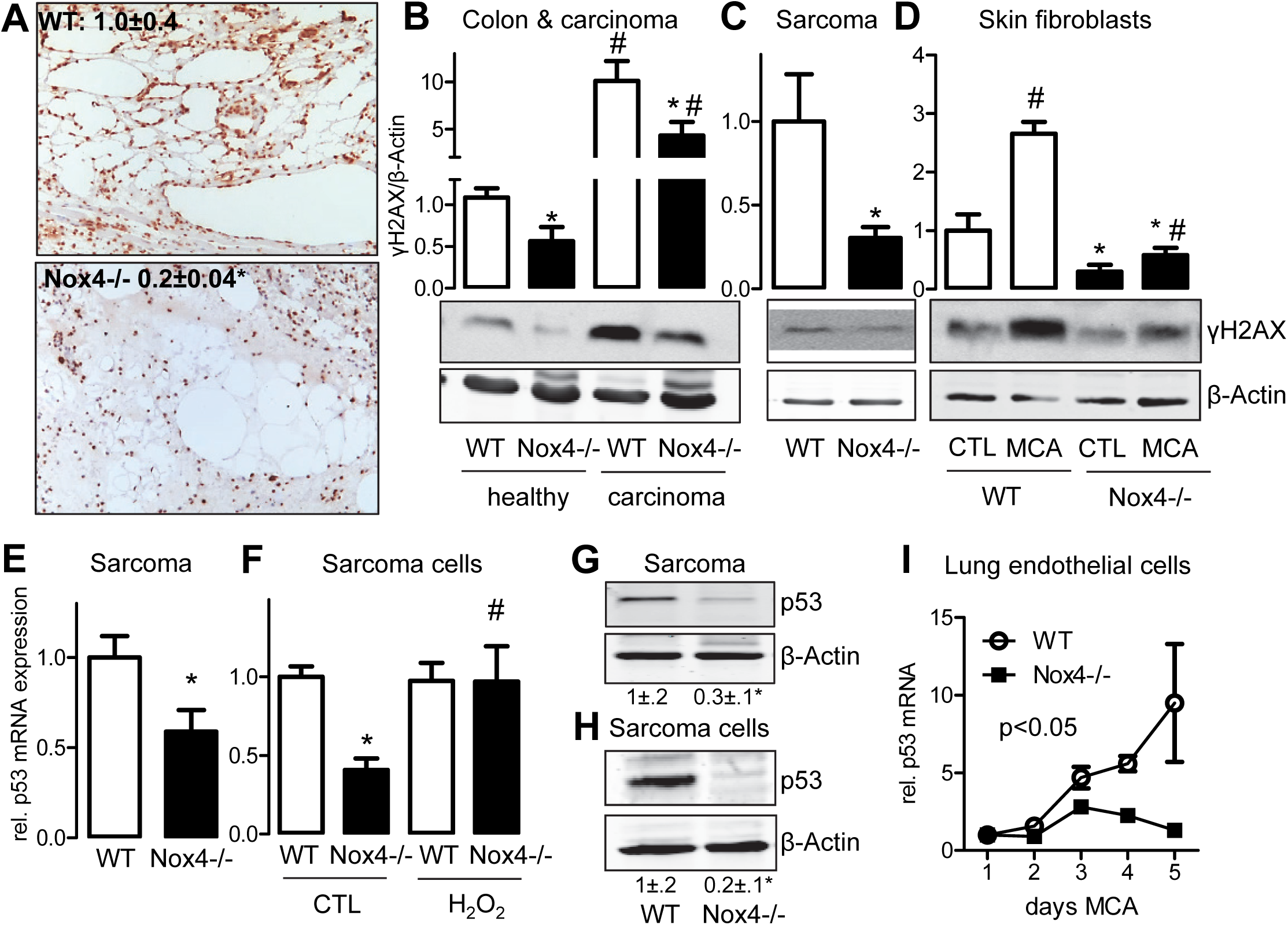
Nox4-deficiency results in an enhanced DNA instability. The expression of the DNA damage marker γH2AX was measured in healthy colon and colon carcinomas (n= 11 WT & 9 Nox4-/-) (**A**) and in fibro sarcoma tissue (**B**) (n=4) by western blot. (**C**) Cross-sections of the skin of WT and Nox4-/- mice stained for phosphorylated H2AX 20 days after MCA injection as a sign for DNA damage. (**D**) Western blot for γH2AX in primary skin fibroblasts treated with or without MCA for 3 days (5μg/ml, n=3). *p<0.05 WT vs. Nox4-/-,#p<0.05 WT/Nox4-/- healthy/CTL vs. WT/Nox4-/- carcinoma/MCA. (**E-H**) p53 abundance. mRNA (**E&F**) and protein (**G&H**) of p53 in fibro sarcoma tissue (**E&G**) and isolated fibro sarcoma cells (**F&H**). Numbers below the blots are the results of the densitometry. *p<0.05. n> 5. H_2_O_2_ denotes treatment with 5μM for 24h. (**I**) p53 mRNA expression was measured by RT-qPCR after MCA treatment (1-5 days, 5μg/ml) in freshly isolated lung endothelial cells (n=5)

### Nox4-deficiency enhances nuclear PP2A activity

Steady state levels of γH2AX depend on the activity of the H2AX phosphorylating ATM and the γH2AX phosphatase. The latter task is carried out by the important serine-threonine phosphatases of the PP2A family (28, 29). Given that ATM expression and activity was not altered after Nox4 knockout, we hypothesized that nuclear PP2A activity is increased in Nox4-/- cells. Indeed, by proximity ligation assay (PLA) with a pan-C-subunit antibody of the PP2A family an increased association of a PP2A member with γH2AX in the nucleus was observed (Fig. 3A). Whereas the total cellular expression of PP2A was similar between WT and Nox4-/- cells, the nuclear PP2A abundance was greatly increased in Nox4-/- cells as judged by the abundance of the C-subunit (Fig. 3B). As a consequence, global serine-threonine phosphatase activity was decreased in the cytosol and increased in the nucleus of Nox4-/- cells when compared to the corresponding WT cells (Fig. 3C). In order to link these data to PP2A activity, experiments with the pan-PP2A inhibitor okadaic acid were carried out at a concentration of 1 nM. At this concentration, the inhibitor is thought to be very selective for PP2A, whereas at higher concentrations, other phosphatases are blocked too. (30). While okadaic acid (1 nM) had no significant effect on nuclear phosphatase activity in WT cells, it reduced the phosphatase activity in Nox4-/- cells to the level observed in WT cells (Fig. 3C). In keeping with this, okadaic acid restored the level of γH2AX in Nox4-/- cells to the level of WT cells but had no effect on γH2AX level in WT cells (Fig. 3D). Strikingly, H_2_O_2_, the product of Nox4 also restored γH2AX levels (Fig. 3E) and reduced the nuclear PP2A level in Nox4-/- cells to the level of WT cells (Fig. 3F). Collectively, these observations suggest that Nox4-derived H_2_O_2_ facilitates accumulation of PP2A in the cytosol. In the absence of Nox4, and thus lack of H_2_O_2_, PP2A accumulates in the nucleus where it dephosphorylates γ2HAX. The result is an attenuated DNA damage response and genomic instability.

**Figure 3.**
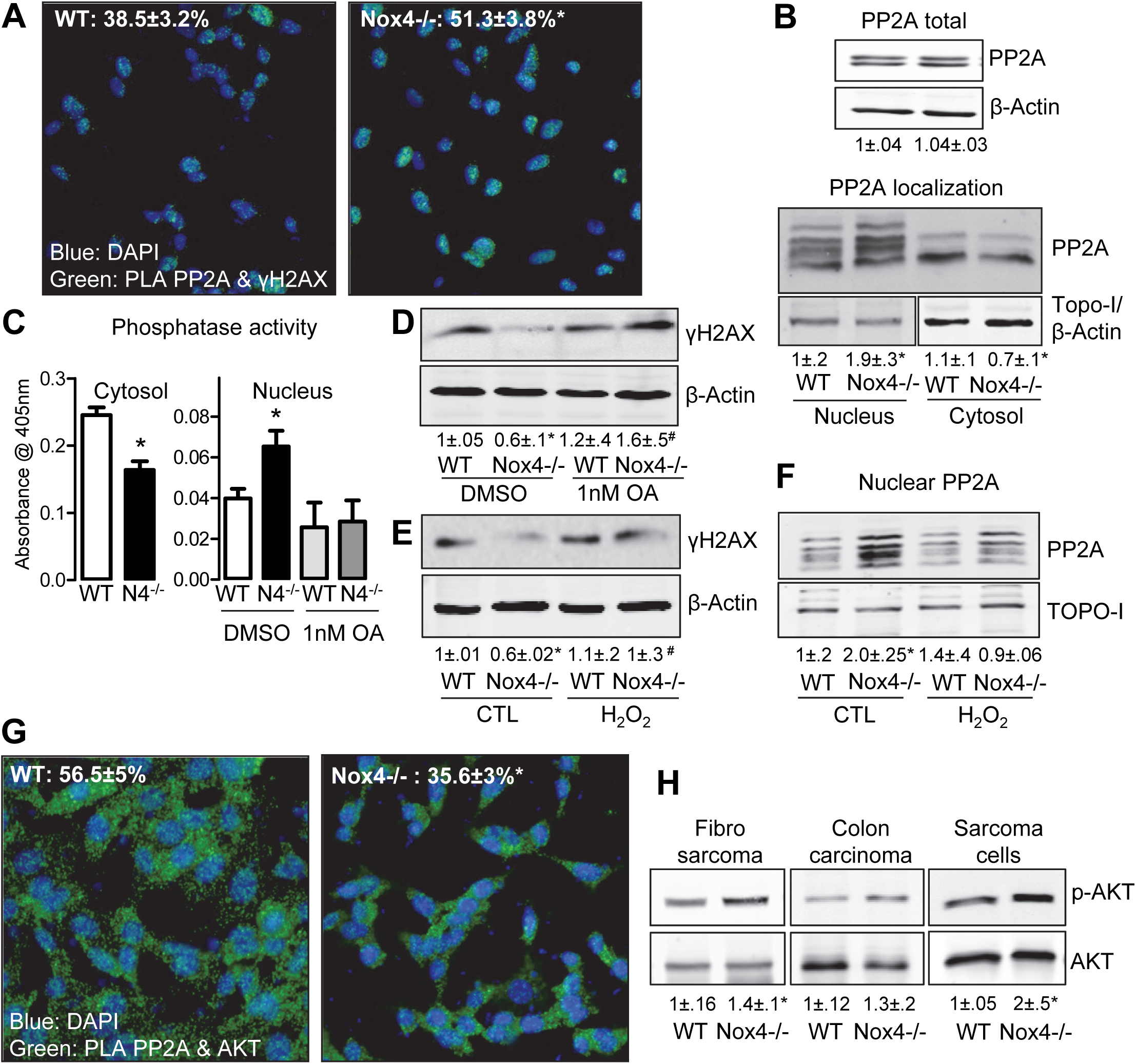
Nox4-deficiency enhances nuclear PP2A activity, this decreases nuclear γH2AX but increases cytosolic AKT phosphorylation. Interaction between PP2A and γH2AX as determined by proximity ligation assay. Quantification of co-localization relative to the number of cells stained with DAPI in %. (**B**) PP2A abundance in cytosolic and nucleus fraction as evaluated by western blot. (**C**) Serine-threonine phosphatase activity as measured under basal conditions and after treatment with okadaic acid (OA) in cytosol and nucleus with the aid of NPP as an artificial substrate. (**D&E**) γH2AX abundance as quantified by immunoblotting with and without 1nM of the PP2A inhibitor okadaic acid (OA) overnight (**D**) or H_2_O_2_ (24h, 5 μM, **E**). (**F**) Nuclear expression of PP2A after treatment with or without H_2_O_2_ as determined by western blot. All experiments were carried out in isolated fibrosarcoma cells (**F**). Statistics are represented as numbers below the representative western blots; *p<0.05 WT vs. Nox4-/-,#p<0.05 Nox4-/- CTL vs. Nox4-/- OA/H_2_O_2_, n=5. (**G**) Interaction between AKT and PP2A as determined by proximity ligation assay. Quantification of co-localization relative to the number of cells stained with DAPI in % (**H**). Western blot for phosphorylation of AKT at Ser473 in fibro sarcoma tissue, colon carcinoma tissue and isolated fibro sarcoma cells of WT and Nox4 -/- mice.

### Nox4 retains PP2A in the cytosol where it interacts with and dephosphorylates AKT

The identification of the molecular basis of the nuclear PP2A accumulation after Nox4 knockout is a complex problem, given the heterogeneous composition of PP2A family members and the fact that numerous mechanisms could account for the effect. Among them are post-translational modifications of the target protein leading to altered localization and stability, changes in the activity of the nuclear import and export machinery and protein-protein interactions which could retain the protein in a compartment. Given the sophisticated regulation of the PP2A enzyme complexes and their numerous targets, literature mining was performed to obtain a plausible candidate mechanism. Of the cytosolic proteins promoting tumor survival, the family of AKT kinases is of outstanding importance (31). AKT is expressed in high abundance in tumors (32) and PP2A family members are responsible for the dephosphorylation of AKT (33, 34). Importantly, in a very different context it was suggested that AKT can be oxidized which promotes its interaction with PP2A (35). On this basis we speculate that the cytosolic interaction of PP2A and AKT could contribute to the mechanism of the altered localization of PP2A.

If our working model was correct, nuclear translocation of PP2A in the absence of Nox4 should result in a decreased association of PP2A with its target AKT and a subsequent increase in AKT phosphorylation, which was indeed the case (Fig. 3G&H). Importantly, we found no evidence that the pathway leading to increased AKT phosphorylation, like the formation of PIP3 or PTEN oxidation, expression and activity were responsible for the increase in AKT phosphorylation (Supplemental figure 5&6).

### AKT is a redox-target of Nox4

In order to determine whether AKT could be the redox-switch responsible for PP2A translocation, Redox-BIAM switch assays were performed, which indeed demonstrated that loss of Nox4 resulted in a significant reduction of AKT oxidation (Fig. 4A). As targeted redox-proteomics approach did not yield sufficient coverage of the peptide sequence of AKT isolated out of tissues, the redox-active cysteines of AKT were mapped from the recombinant protein. AKT was reduced and subsequently incubated with low amounts of H_2_O_2_. This redox-stimuli resulted in disulfide bridge formation between cysteine 60 and 77 and between cysteine 296 and 310, the latter being conserved in all AKT homologues (Fig. 4B & Supplemental figure 7, **Supplemental table 1&2**). Based on these data it could be speculated that Nox4-dependent H_2_O_2_ formation changes the confirmation of AKT by altering disulfide bridge formation.

**Figure 4.**
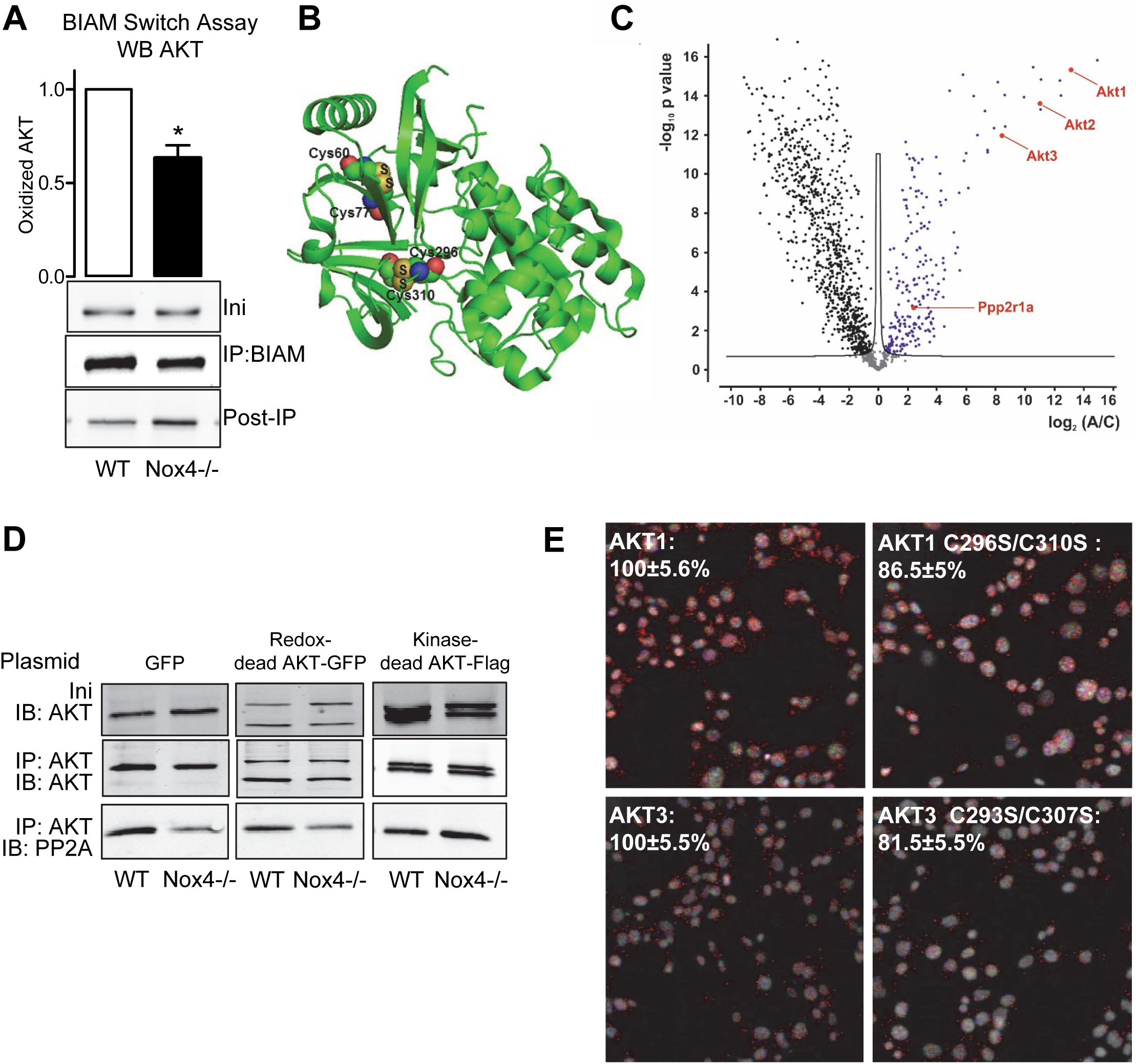
The interaction of PP2A and AKT is redox-sensitive. AKT redox modification as analyzed by BIAM Switch Assay in fibro sarcoma cells of wildtype and Nox4-/- (n=3). (**B**) Mapping of the H_2_O_2_-dependently formed disulfide bridges on the structure of human AKT1 as obtained from 4EJN [25] (see Supplementary table 1&2. (**C**) AKT1-3 proteins were immune-captured and co-purified interacting proteins were identified by LC/MS. Lable free quantification (LFQ) values were statistical analyzed using permutation test (FDR < 0.05, 250 randomizations), see supplementary table 3. Blue and red dots represent significant enriched proteins. Black and grey dots were enriched in negative control or background, respectively. Marked protein Ppp2r1a was found to be significant enriched in AKT1-3 pull-down samples. (**D**) Co-immunoprecipitation of AKT followed by western blotting for PP2A catalytic subunit after overexpression with GFP, the redox-dead mutants for AKT1-3 and phospho-dead AKT1 mutant. (**E**) Proximity ligation in wildtype fibro sarcoma cells after overexpression of AKT1, AKT3, AKT1 and AKT3 redox-dead mutants. Antibodies used for AKT1 were PP2A and HA and for AKT3 PP2A and GFP. Results of cells overexpressed with wildtype AKT were set to 100%.

### Oxidized AKT interacts with the PPP2R1A scaffolding subunit of PP2A

Next, the redox-nature of the interaction of PP2A and AKT was defined. Immunoprecipitation of AKT followed by proteomics analysis revealed that PPP2R1A co-precipitates with AKT (Fig. 4C, **Supplemental table 3**). PPP2R1A, also known as PP2A, subunit A, R1-α isoform or PR65-α, is a scaffolding molecule that coordinates the assembly of the catalytic C subunit and the variable regulatory B subunit. Unfortunately no B or C subunit of PP2A was recovered in these screens. This might be consequence of the fact that the PPP2R1A subunit of PP2A mediates the interaction with other proteins (36). Potentially, detergent conditions were so stringent in order to gain specificity that the PP2A protein complex was disrupted.

To substantiate our findings, an antibody directed against both C-subunit of PP2A was used (37) and regular co-immunoprecipitation experiments followed by Western blot were performed. These experiments demonstrated an interaction of PP2A subunit C with AKT in WT cells, which was impaired in Nox4-/- cells (Fig. 4D) Overexpression of redox dead mutants of AKT (C296S C310S) did not restore co-precipitation of AKT with PP2A in Nox4 knockout cells, whereas after overexpression of a kinase-dead mutant (T308A/S473A) more PP2A was co-precipitated with AKT in the knockout cells (Fig. 4D). In keeping with this, PLA assays demonstrated that the redox-dead mutants exhibited an attenuated interaction with AKT even in Nox4 expressing WT cells (Fig. 4E). Collectively, these data suggest that interaction of PP2A with AKT occurs through the subunit PPP2R1A and requires Nox4-dependent oxidation of AKT but not the kinase function of AKT.

### Kinase-but not Redox-dead AKT restores normal cell function in Nox4-/- cells

Next it was tested whether cytosolic retention of PP2A in the cytosol of WT cells is dependent on the oxidation site of AKT. Overexpression of any of the three wild-type AKT homologues restored cytosolic retention of PP2A. Importantly, this effect was also observed when kinase-dead versions of AKT but not the redox-indicative cysteine-mutant of AKT was overexpressed (Fig. 5A&B). According to our model, restoration of cytosolic trapping of PP2A should also restore DNA damage detection and thus phosphorylation of H2AX. This was indeed the case (Supplemental figure 8). Overexpression of wild-type AKT restored γH2AX in Nox4-/- cells to the level observed in WT cell. In contrast, overexpression of redox-dead AKT failed to have this effect. This would suggest that also DNA-repair is dependent on the oxidation of AKT resulting in PP2A trapping. To address this, comet assays were performed after overexpression of the different AKT plasmids. Wild-type AKT reduced the number of comets in Nox4-/- cells to the level of WT cells and similar effects were observed in kinase-dead mutants of AKT (Fig. 5C&D). In contrast, none of the redox-dead versions of AKT was able to affect the number of comets. Thus, retention of PP2A by Nox4-oxidized AKT stabilized γH2AX in the nucleus, which results in DNA repair and reduced number of comets.

**Figure 5.**
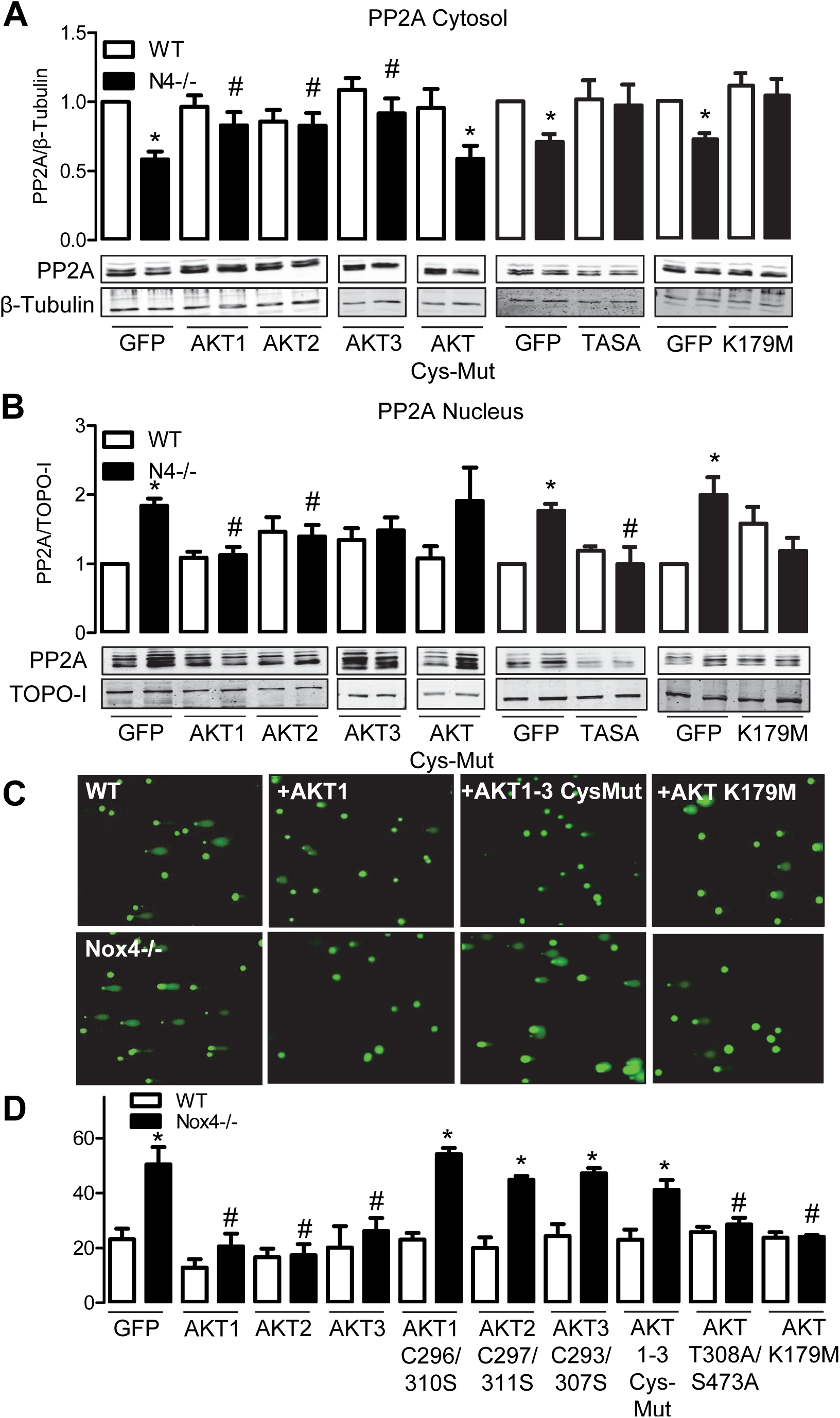
PP2A translocation can be rescued by overexpressing AKT. PP2A (catalytic subunit) localization in the cytosol (**A**) and nucleus (**B**) was determined by western blot after nuclear extraction. Cells were overexpressed with the plasmids indicated. *p<0.05; WT/WT mutated AKT vs. Nox4-/-/Nox4-/- mutated AKT, #p<0.05 Nox4-/- vs. Nox4-/- AKT OE, n=5) (**C**) Comet Assay, Exemplary photos (**C**) and statistics (**D**) of WT and Nox4-/- cells with overexpression of the plasmids indicated *p<0.05 WT vs Nox4-/-, #p<0.05 Nox4 GFP vs. Nox4-/- AKT mutant. n=5

## Discussion

In the present study we observed that genetic deletion of the NADPH oxidase Nox4 resulted in genomic instability, which was a consequence of attenuated DNA damage response due to an increased dephosphorylation of H2AX and attenuated accumulation of p53. Under physiological conditions, Nox4 provides a basal tone of H_2_O_2_ which oxidizes AKT and subsequently sequesters the phosphatase PP2A in the cytosol. Deletion of Nox4 and thus attenuated formation of H_2_O_2_ results in reduced AKT, which does not bind PP2A. As consequence, the phosphatase is transported into the nucleus and dephosphorylates γH2AX. DNA damage accumulates and results in genomic instability and tumor progression.

It has been widely accepted that ROS are potential harmful chemicals which facilitate lipid peroxidation und DNA damage. ROS are mediators of UV and X-ray radiation as well as environmental toxins like arsenide and cigarette smoke. These kinds of damaging ROS are, however, highly reactive, like those forms produced by Fenton or Haber-Weiss chemistry in the presence of transition metals (38, 39). Despite this, redox-reactions are ubiquitous in an oxygen respiring cell and a complex thiol-redox system has developed in aerobic organism to cope with the oxidizing power of oxygen. Cells very well resist a fairly broad concentration range of oxidants and low reactive ROS and in particular H_2_O_2_ have evolved as important signaling molecules. Endogenous ROS generator systems, like the Nox family, utilize antioxidant response systems to mediate signal transduction (40). Moreover, during evolution systems have developed which are redox-sensitive and are transiently inactivated during signaling to allow signal propagation.

Work from several labs including ours suggests that the NADPH oxidase Nox4 has a unique function in this respect. Nox4 constitutively produces H_2_O_2_ and thereby provides a basal oxidative tone to the cell. Through this Nox4 is protective (9) and maintains the activity of antioxidant systems like Nrf2 (10, 41) and protein kinase A (42)(43). H_2_O_2_ produced by Nox4 contributes to differentiation (16) which is in accordance to the concept that during the progression from stem cells towards differentiated cells, cellular ROS production is increased (44). In fact, hypoxia and exquisitely low ROS level are characteristics of stem cells and stem cells niches (44).

Previously, it was noted that deletion of Nox4 results in attenuated differentiation and inflammatory activation which was in part mediated by attenuated eNOS expression, Nrf2 activity (10, 45, 46) and JNK signaling (16). With the present work we establish AKT as an additional redox-target of Nox4 important for malignant transformation. Nox4 promoted the interaction of PP2A with AKT, which facilitated AKT dephosphorylation. Given that AKT promotes survival and proliferation, this finding explains the increased proliferation after deletion of Nox4, observed not only in this but also other studies (39).

As a second effector of Nox4, H2A phosphorylation which is an indicator of DNA damage, was identified. In case of DNA double-strand breaks, H2A is phosphorylated by ATM at Ser139 (47) and this event is a prerequisite to cellular response to DNA-damage (29). The present observation of attenuated γH2AX in the absence of Nox4 is in line with a previous report that antioxidants reduce the level of γH2AX in murine lung cancer tumors (3). It appears that the binding of PP2A to the highly abundant AKT deprives other cellular compartments from this phosphatase. After Nox4 knockout AKT is less oxidized, PP2A is released from AKT and translocates into the nucleus, where it dephosphorylates γH2AX (28).

Phosphatases are well known redox-switches as several families, such as dual-specific phosphatases, the lipid phosphatase PTEN and tyrosine phosphatases are inactivated by H_2_O_2_. In fact, it has been shown that Nox1 inactivates PTEN (48) and inhibition of Nox1 induces apoptosis by attenuating AKT signaling in cancer cells (49). PTP-inactivation by Nox4-derived H_2_O_2_ is certainly occurring under certain conditions and might promote survival of some cancer cells (50, 51); the tonic activity of Nox4, however, in most cells should result in an induction of the antioxidant defense to counteract this process (41, 46, 52).

PP2A, the phosphatase of concern in the present work, however does not contain a redox-sensitive cysteine in the active center but a metal (53), and is therefore not redox-sensitive. The variable intracellular distribution of PP2A and its effect on cell survival might explain the conflicting data obtained with PP2A inhibitors in cancer. There is this long standing history of okadaic acid and other PP2A inhibitors as carcinogens (54), but there is growing controversy regarding the role of PP2A as tumor suppressor. Inactivation of PP2A triggers tumor-selective cell death (55) and the PP2A inhibitors cantharidin or okadaic acid induce apoptosis of pancreatic cancer cells through persistent phosphorylation of IKKα and a sustained activation of the NF-κB pathway (56). Subcellular localization of PP2A activity may be the key to understand the controversial findings. In fact the B subunit determinates the function of the PP2A complex and localization of the phosphatase activity. Out of the four recognized subfamilies; it is plausible, that a B56 complex may prevent DNA damage repair, if not retained in the cytosol, as a B56ϵ containing PP2A holoenzyme has been described to dephosphorylate γH2Ax (57).

An unexpected result of the present study was that Nox4 deletion elevated AKT phosphorylation. It is known since very long that H_2_O_2_, by inhibition of PTEN and activation of tyrosine kinases increases AKT phosphorylation. Moreover, in several tumor cell lines, Nox4, as a source of H_2_O_2_ was suggested to drive AKT phosphorylation (58–62). Nevertheless, in other studies AKT was identified as an inducer of Nox4 expression (63–66). These studies, however, were all performed in transformed stable tumor cell lines. Thus, genomic instability or malignancy induction is not concerned but rather a general aspect of how H_2_O_2_ promotes migration and proliferation. In line with this, any mechanism reducing H_2_O_2_, like inhibition of mitochondria, Nox1, Nox2 or Nox4 usually leads to similar protective response in these studies (50, 67, 68)

Several studies report increased expression of Nox4 in cancer (60, 69). This is not surprising as Nox4 is induced by hypoxia and fibrotic response, which is common in tumors. Thus, Nox4 induction is unlikely to be causal in those studies. Moreover, despite the fact that commercial antibodies are not specific for Nox4 or are not even detecting the enzyme (70), many of the publications identify Nox4 by its molecular weight in Western blots or immuno histology (for example) (60, 71).

Although our work suggests that Nox4 maintains genomic stability, other studies linked Nox enzymes to the opposite effect. p22phox down-regulation contributes to genomic instability in FLT3 expressing leukemia cells (72) and H-Ras transformed cell lines (73). Due to the selection of the individual cell line, an association of this function to a single Nox enzyme is impossible, and also most any homologue has been associated with DNA damage (74).

These variable results stress that tumor research on cell lines should be interpreted with great caution and that the results frequently cannot be transferred to the in vivo situation.

Our study is not the first linking Nox4 to a rather positive protective function in malignant disease. In hepatic cancer, the Nox4 promoter is silenced which appears to promote hepato carcinogenesis in rats (75). This data is even supported in a study looking at Nox expression in hepatocellular carcinoma (76). High Nox1 expression was associated with less favorable outcome, where high Nox4 expression was beneficial (76, 77). Thus, low Nox4 might thereby be a prerequisite for tumor progression. Recently this view has been supported by a more descriptive study where Nox4 siRNA prevented liver cancer in a xenograft-model (78) and in a study of epidermal growth factor receptor inhibitor induced autophagy in head and neck cancer cells in a Nox4 dependent way (79).

Collectively, with the present work, Nox4 was established as an endogenous source of ROS which maintains genomic stability. This work supports clinical data that antioxidants do not protect against cancer initiation and may even be harmful under certain conditions. The dogmatic view of ROS as bona fide harmful molecules should be queried.

## Acknowledgment

We are grateful to Sabine Harenkamp, Maria Walter and Jana Meisterknecht for their excellent technical support of our study. Furthermore, we thank Itamar Goren and Beate Fisslthaler for providing AKT mutant plasmids.

### Conflict of interest

The authors of this manuscript have no conflict of interest in the present study.

### Funding

This work was supported by grants from the Deutsche Forschungsgemeinschaft (DFG) (to KS SCHR1241/1-1 and SFB815/TP1, TP8 &Z1) and the Fraunhofer Gesellschaft (graduate school translational research innovation pharma, TRIP).

---

**Supplemental figure 1:**
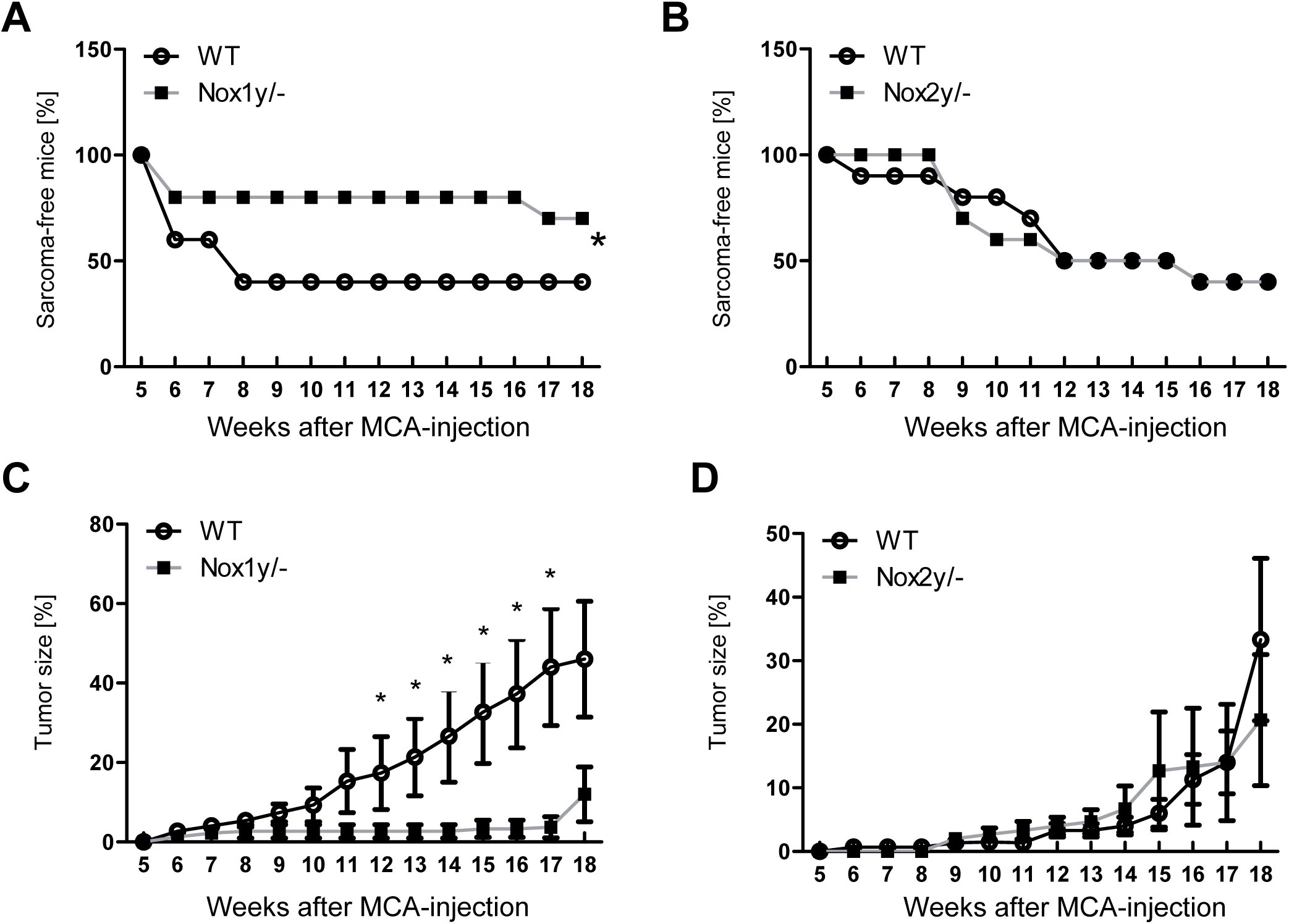
Nox1-deficiency attenuates tumor development whereas the knockout of Nox2 has no effect on tumor development. WT littermates and corresponding Nox1y/- and Nox2y/- mice were injected subcutaneously with MCA. Fibro sarcoma development after MCA-injection in Nox1y/- (**A**) and Nox2y/- mice (**B**) and tumor diameter in % (100% is equal to 1.5 cm) as assessed weekly with a caliper in WT littermates, Nox1y/- (**C**) and Nox2y/- (**D**). *p<0.05 WT vs. Nox1y/-, n=10

**Supplemental figure 2:**
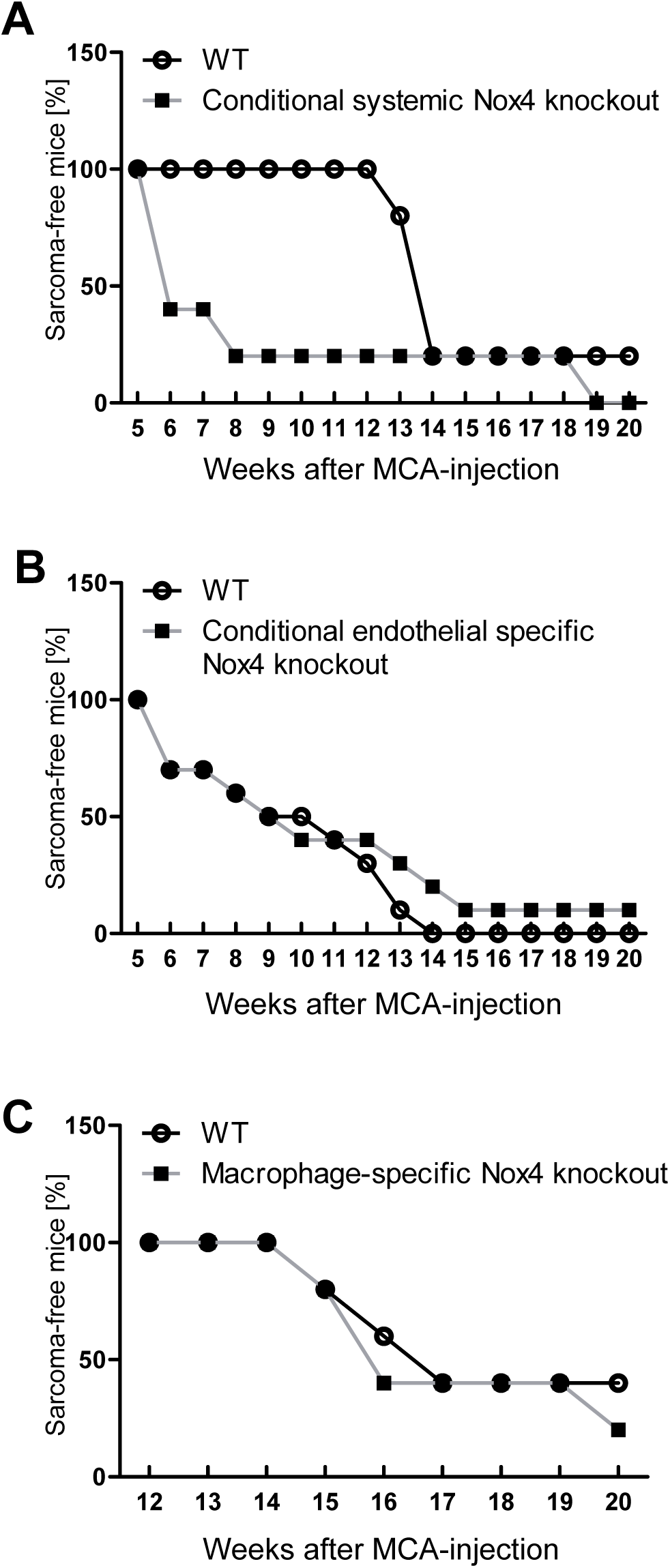
Fibro sarcoma development in response to MCA in conditional Nox4 knockout. (**A**) Nox4^flox/flox^-CMV-CreERT2^+/0^ vs. Nox4^flox/flox^, all after tamoxifen-treatment (**B**) Conditional endothelial-specific (Nox4^flox/flox^-cdh5-CreERT2^+/0^ vs. Nox4^flox/flox^) all after tamoxifen-treatment. (C) LysM-macrophage and LysM neutrophil specific (Nox4^flox/flox^-LysM-Cre^+/0^ vs. Nox4^flox/flox^). Fibro sarcoma development was assessed weekly with a caliper. Mice were sacrificed when tumor reached 1.5 cm, n=4-10

**Supplemental figure 3:**
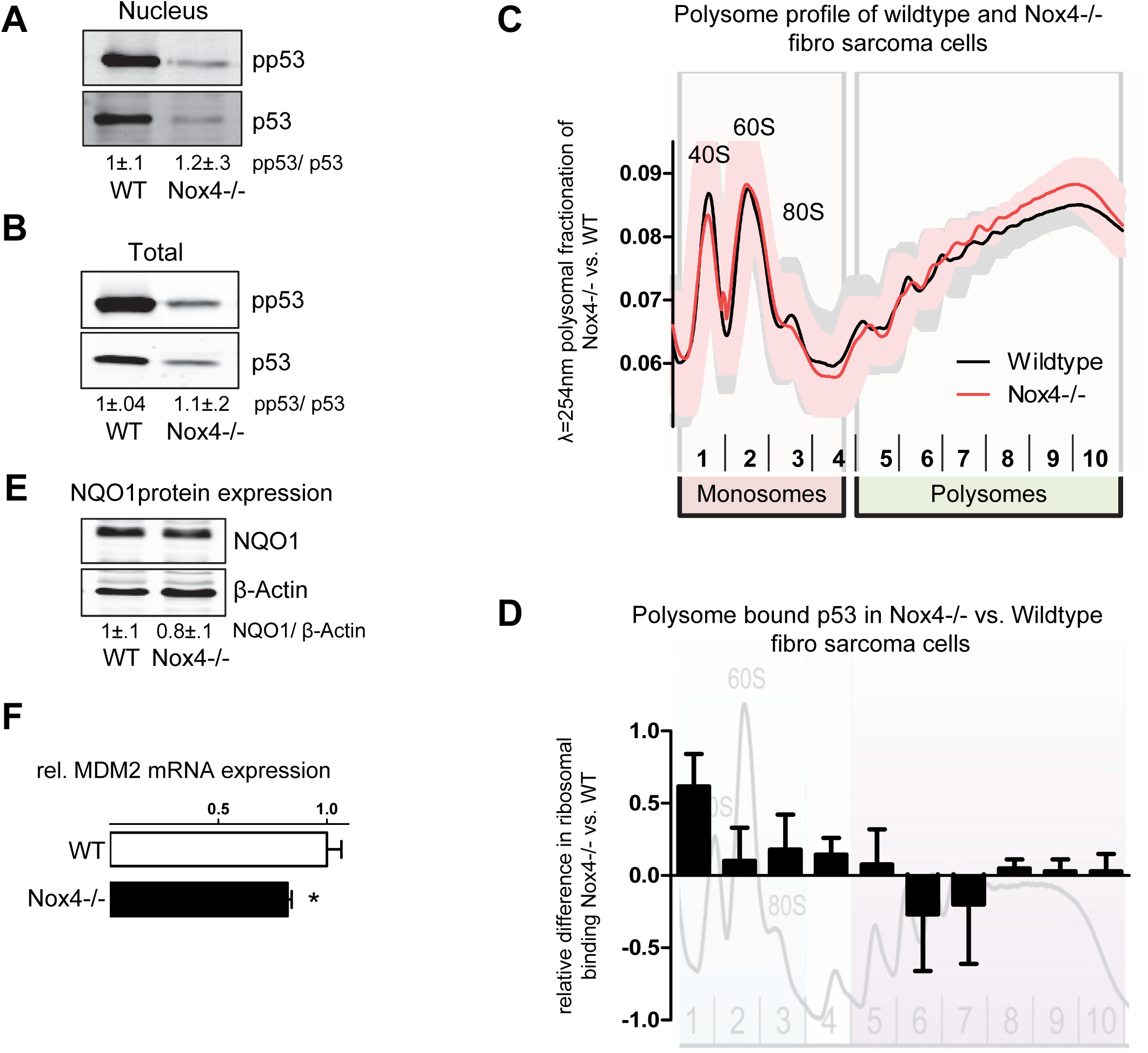
p53 phosphorylation and translation is similar between WT and Nox4-deficient fibro sarcoma cells. (**A&B**) Western blot for p53 and phospho-p53 from nuclear (**A**) and total cellular extracts (**B**) of fibro sarcoma cells. Numbers below the blot indicate the result of the densitometry. (**C&D**) Polysome analysis was performed for investigation of translation. Absorbance was measured at 254 nm after polysomal fractionation in WT and Nox4-/- fibro sarcoma cells (**C**). Fractions were collected and analyzed for p53 mRNA expression with RT-qPCR. Normalization was performed to housekeeping gene GAPDH (**D**). n=3; (**E&F**): NQO1 expression in fibro sarcoma cells was measured by Western blot (**E**). MDM2 (**F**) gene expression as quantified by RT-qPCR. *p<0.05

**Supplemental figure 4:**
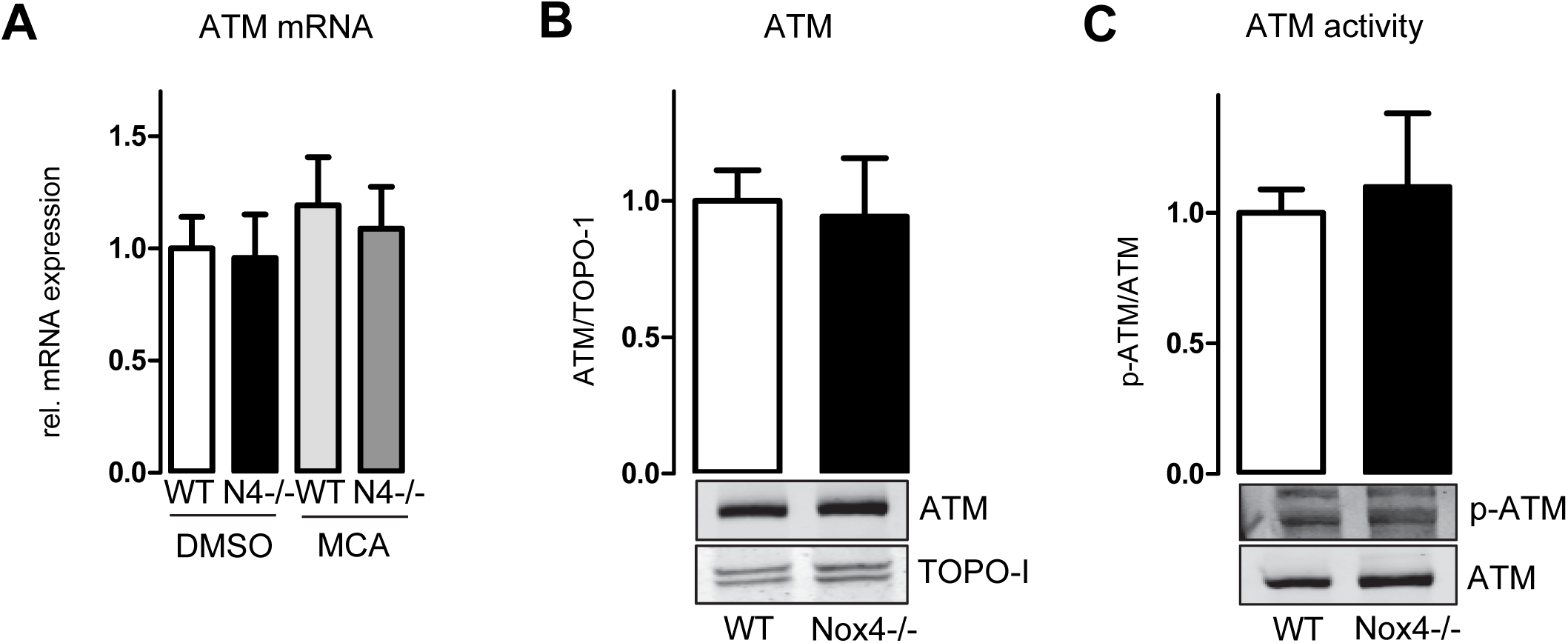
The kinase ATM is not differentially expressed or phosphorylated in Nox4-/-. RT-qPCR was performed in WT and Nox4-/- fibro sarcoma cells after DMSO or MCA treatment for 3 days (5μg/ml). (**B**) ATM expression was quantified by western blot. (**C**) As a readout for its activity the amount of phosphorylated ATM was measured by western blot and normalized to total ATM protein expression. n=6-7

**Supplemental figure 5:**
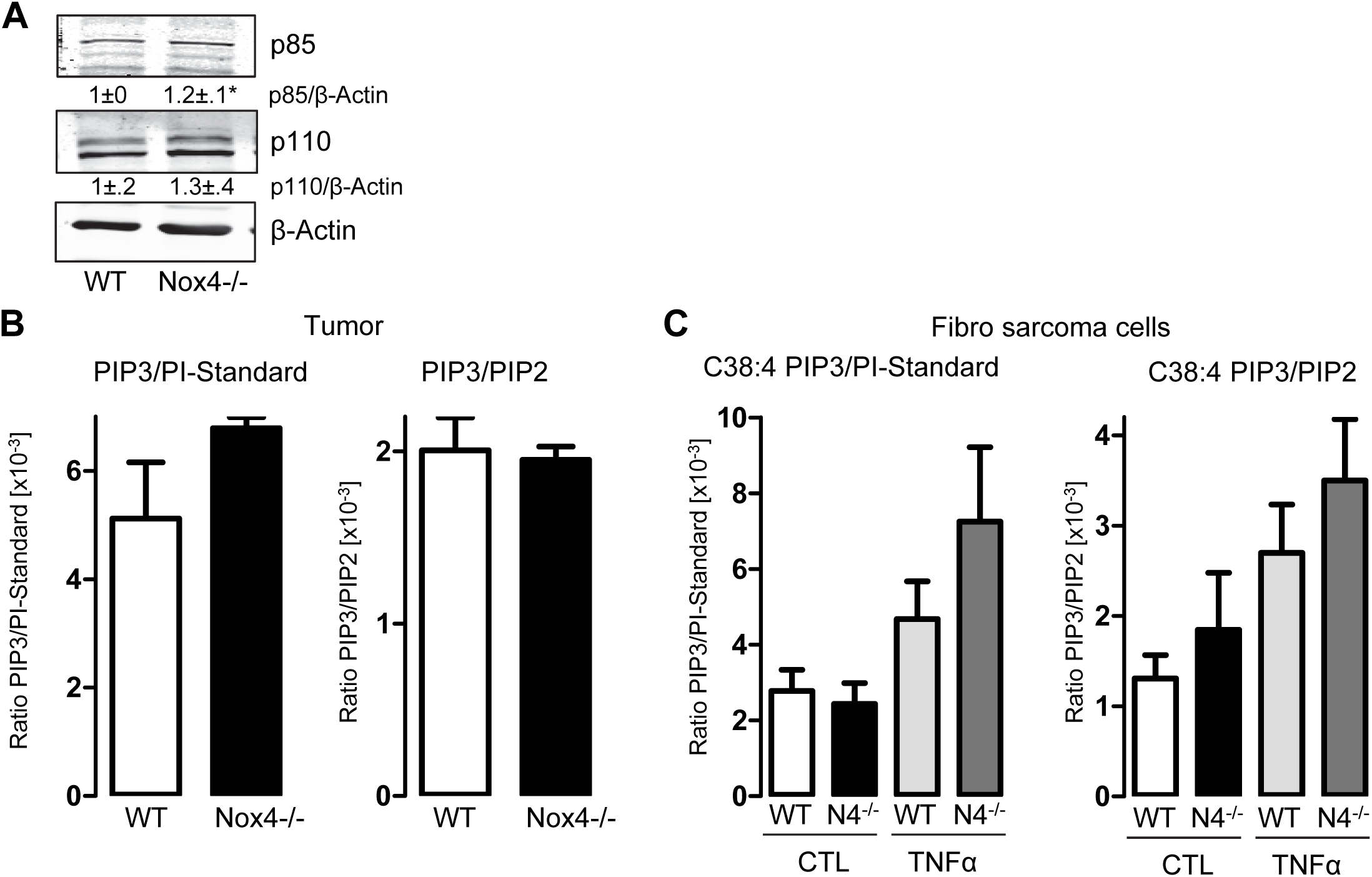
PI3-Kinase activity is unchanged even if expression is slightly increased in the absence of Nox4. Protein abundance (**A**) of the two subunits of the PI3 Kinase, p85 and p110, was assessed by western blot. LC-MS/MS measurements for PIP3 and PIP2 in fibro sarcoma tissue (**B**) and isolated fibro sarcoma cells with or w/o TNFα (10ng/ml) (**C**). Data expressed as C18:0 C20:4 PIP3 response ratio normalized to C18:0 C20:4 PI response ratio to account for cell input variation. Statistics are represented as numbers below the representative western blots; *p<0.05, n=4

**Supplemental figure 6:**
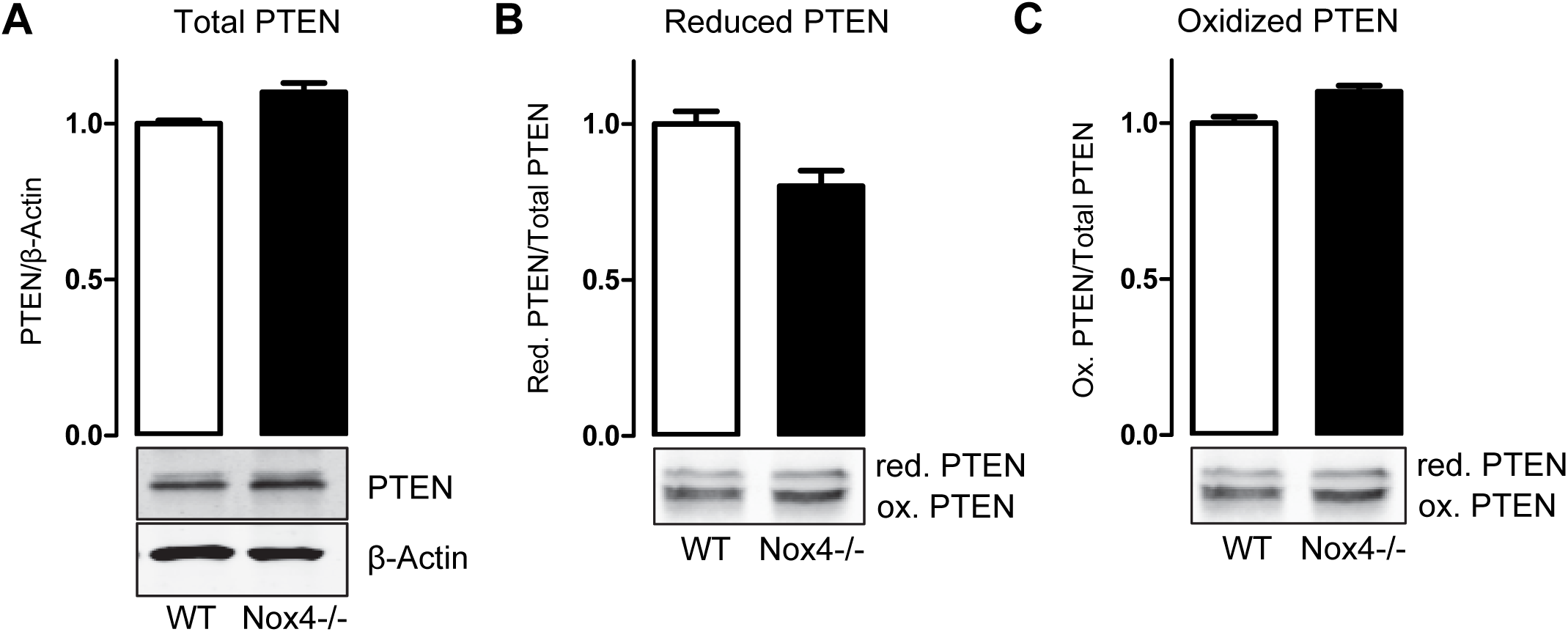
Nox4 knockout in fibro sarcoma cells does not increase the level of reduced PTEN. Total PTEN expression was assessed by western blot. Reduced (**B**) vs. oxidized (**C**) PTEN as evaluated under non-reducing conditions by western blot. n=5

**Supplemental figure 7:**
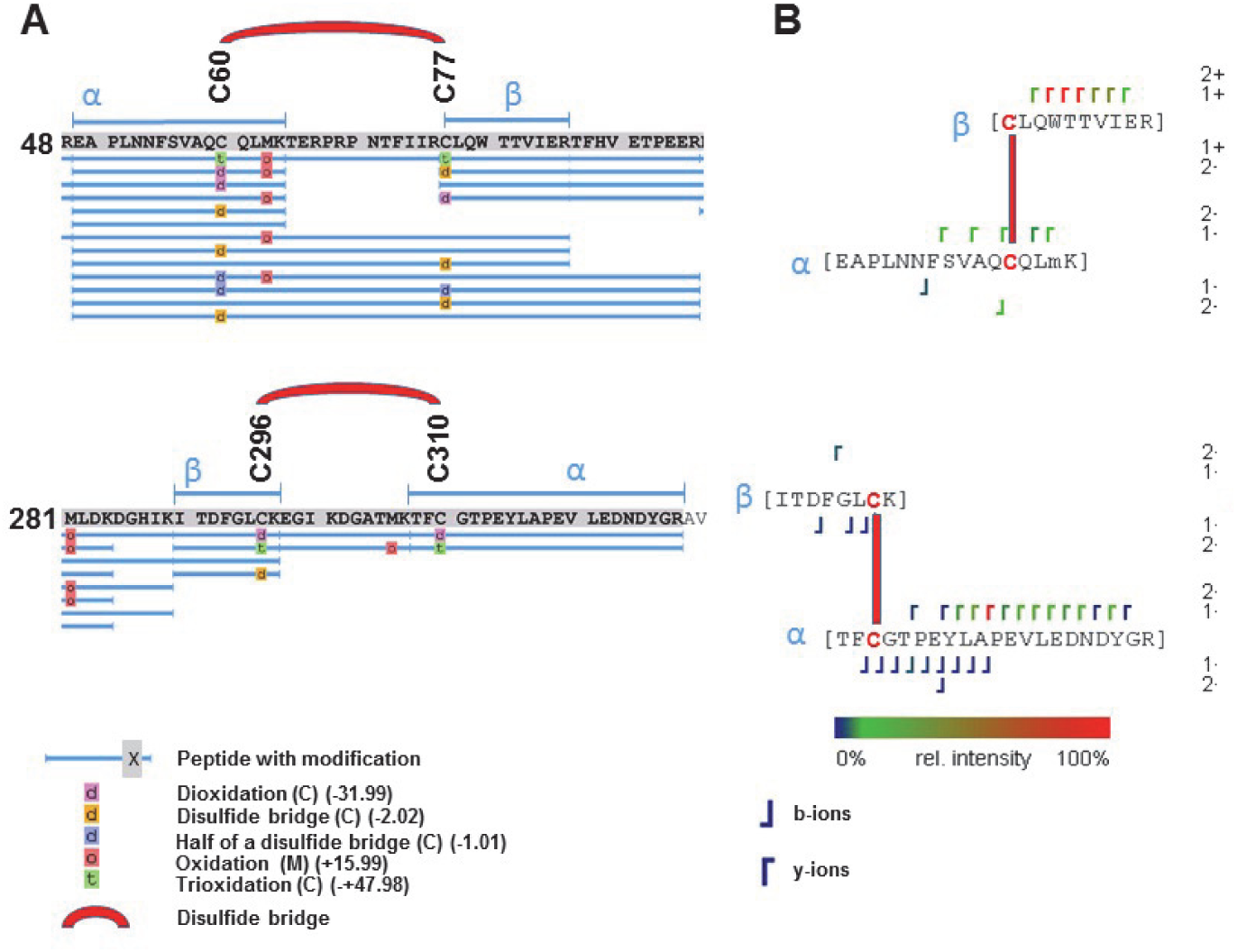
Cys60/77 and Cys296/310 are redox-sensitive and form disulfides. Recombinant human AKT1 (Uniprot ID P31749) was reduced and treated with H_2_O_2_. Samples were digested with trypsin and analyzed by LC/MS. (**A**) The following thiol oxidations on cysteines were identified by proteomics software Peaks7.0: sulfinic acid (Dioxidation), sulfonic acid (Trioxidation) and loss of the mass of one (-1) or two (-2) hydrogens refer to formation of a disulfide bridge. For a list of all identified peptides, see supplementary table 1. (**B**) A special software to identify crosslinked peptides (StavroX, [24]) was further used to identify disulfides. Among the top-scored candidates for disulfides were Cys60/77 and Cys296/310 (Supplementary table 2). Figure shows position of fragment ions within a di-peptide.

**Supplemental figure 8:**
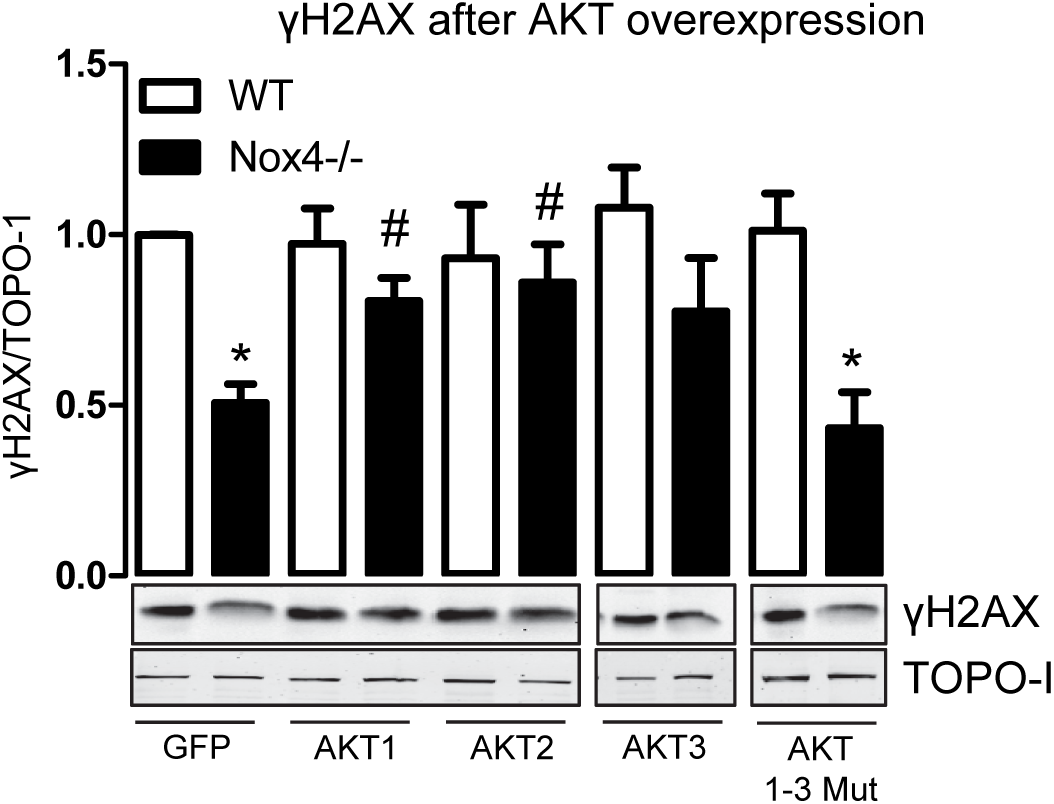
Nox4-dependent oxidation of AKT is necessary for γH2AX phosphorylation and prevention of DNA damage. γH2AX expression determined by western blot after overexpression of the plasmids indicated (n=5); *p<0.05; WT/WT mutated AKT vs. Nox4-/-/Nox4-/- mutated AKT, #p<0.05 Nox4-/- vs. Nox4-/- AKT OE, n=5

## References

[1] Lonn E et al. (2005) Effects of long-term vitamin E supplementation on cardiovascular events and cancer: a randomized controlled trial. JAMA 293:1338–1347.

[2] Omenn GS et al. (1996) Risk factors for lung cancer and for intervention effects in CARET, the Beta-Carotene and Retinol Efficacy Trial. J Natl Cancer Inst 88:1550–1559.

[3] Sayin VI et al. (2014) Antioxidants accelerate lung cancer progression in mice. Sci Transl Med 6:221ra15.

[4] Sirokmany G, Donko A, Geiszt M (2016) Nox/Duox Family of NADPH Oxidases: Lessons from Knockout Mouse Models. Trends in pharmacological sciences 37:318–327.

[5] Brandes RP, Weissmann N, Schröder K (2014) Nox family NADPH oxidases: Molecular mechanisms of activation. Free radical biology & medicine 76C:208–226.

[6] Groeger G, Mackey AM, Pettigrew CA, Bhatt L, Cotter TG (2009) Stress-induced activation of Nox contributes to cell survival signalling via production of hydrogen peroxide. Journal of neurochemistry 109:1544–1554.

[7] Jiang F, Liu G-S, Dusting GJ, Chan EC (2014) NADPH oxidase-dependent redox signaling in TGF-beta-mediated fibrotic responses. Redox biology 2:267–272.

[8] Menegon S, Columbano A, Giordano S (2016) The Dual Roles of NRF2 in Cancer. Trends in molecular medicine 22:578–593.

[9] Schröder K et al. (2012) Nox4 is a protective reactive oxygen species generating vascular NADPH oxidase. Circulation research 110:1217–1225.

[10] Zhang M et al. (2010) NADPH oxidase-4 mediates protection against chronic load-induced stress in mouse hearts by enhancing angiogenesis. Proc Natl Acad Sci U S A 107:18121–18126.

[11] Craige SM et al. (2015) Endothelial NADPH oxidase 4 protects ApoE-/- mice from atherosclerotic lesions. Free radical biology & medicine 89:1–7.

[12] Di Marco E et al. (2016) NOX4-derived reactive oxygen species limit fibrosis and inhibit proliferation of vascular smooth muscle cells in diabetic atherosclerosis. Free radical biology & medicine 97:556–567.

[13] Gray SP et al. (2016) Reactive Oxygen Species Can Provide Atheroprotection via NOX4-Dependent Inhibition of Inflammation and Vascular Remodeling. Arteriosclerosis, thrombosis, and vascular biology 36:295–307.

[14] Clempus RE et al. (2007) Nox4 is required for maintenance of the differentiated vascular smooth muscle cell phenotype. Arterioscler Thromb Vasc Biol 27:42–48.

[15] Schröder K, Wandzioch K, Helmcke I, Brandes RP (2009) Nox4 acts as a switch between differentiation and proliferation in preadipocytes. Arterioscler Thromb Vasc Biol 29:239–245.

[16] Goettsch C et al. (2013) NADPH oxidase 4 limits bone mass by promoting osteoclastogenesis. J Clin Invest 123:4731–4738.

[17] Laleu B et al. (2010) First in class, potent, and orally bioavailable NADPH oxidase isoform 4 (Nox4) inhibitors for the treatment of idiopathic pulmonary fibrosis. J Med Chem 53:7715–7730.

[18] Gavazzi G et al. (2007) NOX1 deficiency protects from aortic dissection in response to angiotensin II. Hypertension 50:189–196.

[19] Bradford MM (1976) A rapid and sensitive method for the quantitation of microgram quantities of protein utilizing the principle of protein-dye binding. Anal Biochem 72:248–254.

[20] Riccardi C, Nicoletti I (2006) Analysis of apoptosis by propidium iodide staining and flow cytometry. Nat Protoc 1:1458–1461.

[21] Rubsamen D et al. (2014) Inflammatory conditions induce IRES-dependent translation of cyp24a1. PloS one 9:e85314.

[22] Clark J et al. (2011) Quantification of PtdInsP3 molecular species in cells and tissues by mass spectrometry. Nat Methods 8:267–272.

[23] Rappsilber J, Mann M, Ishihama Y (2007) Protocol for micro-purification, enrichment, prefractionation and storage of peptides for proteomics using StageTips. Nature protocols 2:1896– 1906.

[24] Gotze M et al. (2012) StavroX--a software for analyzing crosslinked products in protein interaction studies. Journal of the American Society for Mass Spectrometry 23:76–87.

[25] Ashwell MA et al. (2012) Discovery and optimization of a series of 3-(3-phenyl-3H-imidazo4,5-bpyridin-2-yl)pyridin-2-amines: orally bioavailable, selective, and potent ATP-independent Akt inhibitors. Journal of medicinal chemistry 55:5291–5310.

[26] Cox J, Mann M (2008) MaxQuant enables high peptide identification rates, individualized p.p.b.-range mass accuracies and proteome-wide protein quantification. Nature biotechnology 26:1367– 1372.

[27] Tyanova S et al. (2016) The Perseus computational platform for comprehensive analysis of (prote)omics data. Nature methods 13:731–740.

[28] Chowdhury D et al. (2005) gamma-H2AX dephosphorylation by protein phosphatase 2A facilitates DNA double-strand break repair. Mol Cell 20:801–809.

[29] Keogh M-C et al. (2006) A phosphatase complex that dephosphorylates gammaH2AX regulates DNA damage checkpoint recovery. Nature 439:497–501.

[30] Cohen P, Klumpp S, Schelling DL (1989) An improved procedure for identifying and quantitating protein phosphatases in mammalian tissues. FEBS letters 250:596–600.

[31] Amaravadi R, Thompson CB (2005) The survival kinases Akt and Pim as potential pharmacological targets. The Journal of clinical investigation 115:2618–2624.

[32] Bellacosa A, Kumar CC, Di Cristofano A, Testa JR (2005) Activation of AKT kinases in cancer: implications for therapeutic targeting. Advances in cancer research 94:29–86.

[33] Millward TA, Zolnierowicz S, Hemmings BA (1999) Regulation of protein kinase cascades by protein phosphatase 2A. Trends in biochemical sciences 24:186–191.

[34] Kuo Y-C et al. (2008) Regulation of phosphorylation of Thr-308 of Akt, cell proliferation, and survival by the B55alpha regulatory subunit targeting of the protein phosphatase 2A holoenzyme to Akt. The Journal of biological chemistry 283:1882–1892.

[35] Murata H et al. (2003) Glutaredoxin exerts an antiapoptotic effect by regulating the redox state of Akt. J Biol Chem 278:50226–50233.

[36] Krauss G, ed (2014) Biochemistry of signal transduction and regulation (Wiley-VCH, Weinheim).

[37] Kiely M, Kiely PA (2015) PP2A: The Wolf in Sheep’s Clothing? Cancers 7:648–669.

[38] Mello Filho, A C, Hoffmann ME, Meneghini R (1984) Cell killing and DNA damage by hydrogen peroxide are mediated by intracellular iron. Biochem J 218:273–275.

[39] Franke TF, Kaplan DR, Cantley LC (1997) PI3K: downstream AKTion blocks apoptosis. Cell 88:435–437.

[40] Brandes RP, Weissmann N, Schröder K (2014) Redox-mediated signal transduction by cardiovascular Nox NADPH oxidases. Journal of molecular and cellular cardiology 73:70–79.

[41] Brewer AC et al. (2011) Nox4 regulates Nrf2 and glutathione redox in cardiomyocytes in vivo. Free radical biology & medicine 51:205–215.

[42] Burgoyne JR et al. (2015) Deficient angiogenesis in redox-dead Cys17Ser PKARIalpha knock-in mice. Nature communications 6:7920.

[43] Burgoyne JR, Mongue-Din H, Eaton P, Shah AM (2012) Redox signaling in cardiac physiology and pathology. Circulation research 111:1091–1106.

[44] Ushio-Fukai M, Rehman J (2014) Redox and metabolic regulation of stem/progenitor cells and their niche. Antioxidants & redox signaling 21:1587–1590.

[45] Goy C et al. (2014) The imbalanced redox status in senescent endothelial cells is due to dysregulated Thioredoxin-1 and NADPH oxidase 4. Mitochondria, Metabolic Regulation and the Biology of Aging 56:45–52.

[46] Smyrnias I et al. (2014) Nicotinamide Adenine Dinucleotide Phosphate Oxidase-4-Dependent Upregulation of Nuclear Factor Erythroid-Derived 2-Like 2 Protects the Heart During Chronic Pressure Overload. Hypertension.

[47] Bonner WM et al. (2008) GammaH2AX and cancer. Nature reviews. Cancer 8:957–967.

[48] Cui W et al. (2011) NOX1/nicotinamide adenine dinucleotide phosphate, reduced form (NADPH) oxidase promotes proliferation of stellate cells and aggravates liver fibrosis induced by bile duct ligation. Hepatology 54:949–958.

[49] Ito K et al. (2016) Inhibition of Nox1 induces apoptosis by attenuating the AKT signaling pathway in oral squamous cell carcinoma cell lines. Oncology reports 36:2991–2998.

[50] Jayavelu AK et al. (2016) NOX4-driven ROS formation mediates PTP inactivation and cell transformation in FLT3ITD-positive AML cells. Leukemia 30:473–483.

[51] Mondol AS, Tonks NK, Kamata T (2014) Nox4 redox regulation of PTP1B contributes to the proliferation and migration of glioblastoma cells by modulating tyrosine phosphorylation of coronin-1C. Free radical biology & medicine 67:285–291.

[52] Nlandu-Khodo S et al. (2016) NADPH oxidase 4 deficiency increases tubular cell death during acute ischemic reperfusion injury. Scientific reports 6:38598.

[53] Guo F et al. (2014) Structural basis of PP2A activation by PTPA, an ATP-dependent activation chaperone. Cell Res. 24:190–203.

[54] Fujiki H, Suganuma M (1993) Tumor promotion by inhibitors of protein phosphatases 1 and 2A: the okadaic acid class of compounds. Adv Cancer Res 61:143–194.

[55] Zimmerman R et al. (2012) PP2A inactivation is a crucial step in triggering apoptin-induced tumor-selective cell killing. Cell Death Dis 3:e291.

[56] Li W et al. (2011) PP2A inhibitors induce apoptosis in pancreatic cancer cell line PANC-1 through persistent phosphorylation of IKKalpha and sustained activation of the NF-kappaB pathway. Cancer Lett 304:117–127.

[57] Li X, Nan A, Xiao Y, Chen Y, Lai Y (2015) PP2A-B56 complex is involved in dephosphorylation of gamma-H2AX in the repair process of CPT-induced DNA double-strand breaks. Toxicology 331:57–65.

[58] Tanaka M et al. (2015) Inhibition of NADPH oxidase 4 induces apoptosis in malignant mesothelioma: Role of reactive oxygen species. Oncology reports 34:1726–1732.

[59] Zhao QD et al. (2015) NADPH oxidase 4 induces cardiac fibrosis and hypertrophy through activating Akt/mTOR and NFkappaB signaling pathways. Circulation 131:643–655.

[60] Zhang C et al. (2014) NOX4 promotes non-small cell lung cancer cell proliferation and metastasis through positive feedback regulation of PI3K/Akt signaling. Oncotarget 5:4392–4405.

[61] Jafari N et al. (2017) CRISPR-Cas9 Mediated NOX4 Knockout Inhibits Cell Proliferation and Invasion in HeLa Cells. PloS one 12:e0170327.

[62] Crosas-Molist E et al. (2016) The NADPH oxidase NOX4 represses epithelial to amoeboid transition and efficient tumour dissemination. Oncogene.

[63] Liu Z-M, Tseng H-Y, Tsai H-W, Su F-C, Huang H-S (2015) Transforming growth factor beta-interacting factor-induced malignant progression of hepatocellular carcinoma cells depends on superoxide production from Nox4. Free radical biology & medicine 84:54–64.

[64] Choi J-A, Jung YS, Kim JY, Kim HM, Lim IK (2016) Inhibition of breast cancer invasion by TIS21/BTG2/Pc3-Akt1-Sp1-Nox4 pathway targeting actin nucleators, mDia genes. Oncogene 35:83–93.

[65] Edderkaoui M et al. (2013) NADPH oxidase activation in pancreatic cancer cells is mediated through Akt-dependent up-regulation of p22phox. The Journal of biological chemistry 288:36259.

[66] Govindarajan B et al. (2007) Overexpression of Akt converts radial growth melanoma to vertical growth melanoma. The Journal of clinical investigation 117:719–729.

[67] Zhang H-S, Sang W-W, Ruan Z, Wang Y-O (2011) Akt/Nox2/NF-kappaB signaling pathway is involved in Tat-induced HIV-1 long terminal repeat (LTR) transactivation. Archives of biochemistry and biophysics 505:266–272.

[68] Kodama R et al. (2013) ROS-generating oxidases Nox1 and Nox4 contribute to oncogenic Ras-induced premature senescence. Genes to cells : devoted to molecular & cellular mechanisms 18:32–41.

[69] Roy K et al. (2015) NADPH oxidases and cancer. Clinical science (London, England : 1979) 128:863–875.

[70] Babelova A et al. (2012) Role of Nox4 in murine models of kidney disease. Free radical biology & medicine 53:842–853.

[71] Singh L et al. (2016) Prognostic significance of NADPH oxidase-4 as an indicator of reactive oxygen species stress in human retinoblastoma. International journal of clinical oncology 21:651– 657.

[72] Stanicka J, Russell EG, Woolley JF, Cotter TG (2015) NADPH oxidase-generated hydrogen peroxide induces DNA damage in mutant FLT3-expressing leukemia cells. The Journal of biological chemistry 290:9348–9361.

[73] Weyemi U et al. (2012) ROS-generating NADPH oxidase NOX4 is a critical mediator in oncogenic H-Ras-induced DNA damage and subsequent senescence. Oncogene 31:1117–1129.

[74] Kang MA, So E-Y, Simons AL, Spitz DR, Ouchi T (2012) DNA damage induces reactive oxygen species generation through the H2AX-Nox1/Rac1 pathway. Cell death & disease 3:e249.

[75] Lopez-Alvarez GS et al. (2017) Gene silencing of Nox4 by CpG island methylation during hepatocarcinogenesis in rats. Biology open 6:59–70.

[76] Lu C-L et al. (2011) NADPH oxidase DUOX1 and DUOX2 but not NOX4 are independent predictors in hepatocellular carcinoma after hepatectomy. Tumour biology : the journal of the International Society for Oncodevelopmental Biology and Medicine 32:1173–1182.

[77] Ha SY et al. (2016) NADPH Oxidase 1 and NADPH Oxidase 4 Have Opposite Prognostic Effects for Patients with Hepatocellular Carcinoma after Hepatectomy. Gut and liver 10:826–835.

[78] Crosas-Molist E et al. (2014) The NADPH oxidase NOX4 inhibits hepatocyte proliferation and liver cancer progression. Free radical biology & medicine 69:338–347.

[79] Sobhakumari A et al. (2013) NOX4 mediates cytoprotective autophagy induced by the EGFR inhibitor erlotinib in head and neck cancer cells. Toxicol Appl Pharmacol 272:736–745.

